# An amygdalopontine pathway promotes motor programs of ingestion

**DOI:** 10.1101/2025.06.05.657686

**Authors:** Danielle S. Lafferty, Jeremiah Isaac, Joelyz S. Wolcott, Amy Phan, Lily Reck, Andrew Lutas

## Abstract

Despite internal cues that signal fullness, animals can continue eating when motivated by context or palatability. The neural pathways and signals that enable animals to override these fullness cues remain unclear. We examined a central amygdala (CeA) projection to the dorsolateral pons that targets the parabrachial nucleus, a well-established meal termination center, and the adjacent supratrigeminal nucleus, a region that controls orofacial movements. Activity in this CeA^pons^ pathway correlated with the animal’s licking behavior but was not modulated by metabolic need or palatability cues. CeA^pons^ stimulation caused animals to overeat, consume non-edible objects within reach, or exhibit ingestion-like behaviors—licking, chewing, and grasping—even when no target was present. Depending on training and context, stimulation elicited either licking or pellet consumption, suggesting that CeA^pons^ promotes a flexible, goal-directed ingestive state by recruiting consummatory motor circuits rather than simply suppressing satiety signals. These findings highlight how forebrain-brainstem interactions can re-engage feeding behavior beyond homeostatic need.

## INTRODUCTION

Feeding is normally constrained by internal signals of satiation and satiety, yet animals can override or bypass these controls and continue eating^1^. This behavioral flexibility is evident not only in unique situations like competitive eating^2^ or during stress and emotional distress^3–5^, but also in everyday contexts where palatable foods are consumed beyond metabolic need. These departures from homeostatic regulation suggest that the brain can suppress or bypass circuits that normally inhibit feeding^1,6^. Constraints on feeding are mediated by hindbrain circuits, particularly the nucleus of the solitary tract (NTS) and the parabrachial nucleus (PBN), which integrate interoceptive signals and help terminate ingestion^6–8^. The PBN, located within the dorsolateral pons, is especially important for limiting meal size and suppressing food intake during illness or pain^9–11^. Despite this robust anorexigenic circuitry, animals may overcome it through forebrain modulation. One candidate for this top-down control is the amygdala.

The central nucleus of the amygdala (CeA) contains inhibitory neurons that project to the NTS and PBN. Inhibition of these hindbrain circuits by the CeA may enable animals to override satiety signals and continue ingesting food. Indeed, activation of CeA projections to the PBN (CeA^PBN^) drove overconsumption of food, even in the presence of satiety cues, by potentially enhancing the reinforcing properties of palatable food^12–15^. However, the CeA has also been directly linked to the control of jaw movements such as biting potentially via disinhibition of the orofacial motor nuclei^16,17^. Indeed, studies spanning from the 1950’s to the present have observed a direct effect on orofacial movements by CeA activation^12,15,16,18–22^ and found neural correlates of these movements in recordings of amygdala neurons^23,24^. However, this orofacial motor control by the CeA has not been directly implicated in hedonic feeding. Thus, it remains unclear if the ability of CeA activity to cause overeating is a result of suppressing satiety or promoting ingestive motor programs.

To distinguish between these possibilities, we focused on the CeA^PBN^ projection and adjacent regions in the dorsolateral pons (CeA^pons^). These regions adjacent to the PBN contain premotor circuits controlling orofacial movements which also receive direct input from the CeA^25–29^, but which are not often highlighted in feeding studies. A region just ventral to the PBN called the supratrigeminal area coordinates rhythmic actions such as licking, biting, and chewing via its direct connections with downstream orofacial motor nuclei^30–32^. These circuits generate the patterned outputs required for ingestion and are modulated by input from forebrain regions including the motor cortex, superior colliculus, and amygdala^30^. Through these pathways, higher-order brain areas can shape the initiation and execution of feeding behaviors, enabling animals to adjust motor output to environmental and internal states.

Here, we investigated the CeA^pons^ pathway using fiber photometry, two-photon imaging, and optogenetics. Our results challenge the idea that CeA^PBN^ signaling alone drives overconsumption, as we observed neural activity that was linked to motor actions regardless of what was being ingested and optogenetic activation of this pathways drove rapid orofacial behaviors that caused ingestion of any liquid or solid food as well as nearby non-food objects. Consumption of non-food items and our observations of putative eating-associated behaviors in the absence of any food show that within the pons, CeA-driven ingestion is not facilitated solely by suppressing satiety signals, but more likely by engaging pontine premotor circuits to promote consummatory behavior.

## RESULTS

### CeA^pons^ axon activity increases during bouts of licking and is correlated with bout duration

To determine whether CeA^pons^ axons are active preceding or during ingestive behaviors, we first sought to record the timing of CeA^pons^ axon activity using fiber photometry. We expressed an axon-targeted calcium sensor (axon-GCaMP6s) in GABAergic neurons of the CeA, preferentially aiming for the anterior CeM (Figure 1A). We observed labeled CeA axons throughout the PBN and nearby supratrigeminal and peritrigeminal areas (Figure 1A and Figures S1A-C). Since the photometry signals likely arose from axons in all these regions, we refer to them as CeA^pons^ axons.

**Figure 1:**
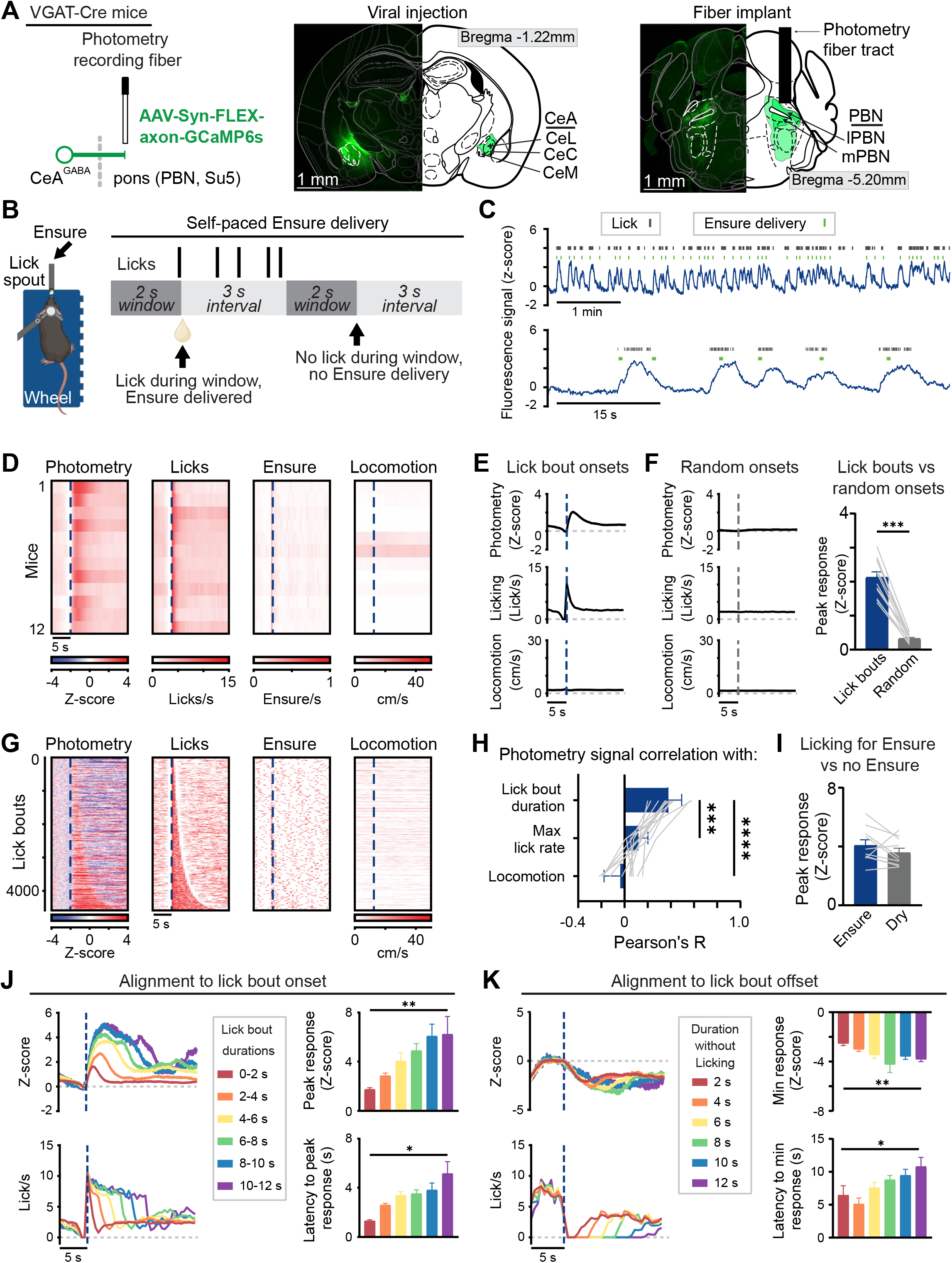
Fiber photometry recordings of calcium signals in CeA^pons^ axons reveals increases in activity following onset of bouts of licking that are correlated with licking duration. (A) Left: schematic depicting fiber photometry recording of GCaMP6s signals in CeA axons in the PBN. Middle: representative image of GCaMP6s expression in CeA. Right: representative GCaMP6s labeled axons in the PBN and other pontine regions with optical implant above. (B) Left: schematic depicting head fixation on a treadmill and available lick spout delivering Ensure. Right: schematic of the task structure in which licking triggered Ensure delivery. (C) Representative traces of GCaMP6s fluorescence showing signal dynamics around the timing of licking and Ensure delivery. (D) Heatmap showing the average response aligned to the onset of lick bouts of individual mice. (E) Time course of mean GCaMP6s photometry, licking, and locomotion aligned to lick bout onset. (F) Left: time course of the mean photometry, licking, and locomotion aligned to random onsets. Right: Peak GCaMP6s response following lick bout or random onsets. Two-tailed paired t test. (G) Heatmap showing signals during every lick bout across all mice with rows sorted by the duration of the lick bout. (H) Pearson’s correlation coefficient calculated for each individual mouse comparing photometry area under the curve (AUC) with lick bout duration, maximum lick rate during the lick bout, or locomotion AUC. Repeated-measures one-way ANOVA with Dunnett’s correction for multiple comparisons. (I) Peak response for lick bouts either containing or lacking Ensure. Bouts have been matched to equalize the distribution of licking behavior. Two-tailed paired t test. (J) Left: time courses of mean photometry for lick bouts of differing durations aligned to lick bout onsets. Right: peak response and latency to peak for these binned lick bouts. Mixed-effects analysis with Tukey’s correction for multiple comparisons. (K) Left: time courses of mean photometry aligned to lick bout offsets and binned by the duration without licking following the offset. Right: minimum response and latency to minimum for these binned offset signals. Mixed-effects analysis with Tukey’s correction for multiple comparisons. All data: N = 12 mice (7M, 5F), mean ± SEM.

To examine the precise timing of behavior during ingestion, we used an apparatus in which head-fixed mice could locomote on a wheel while obtaining liquid food (Ensure^®^ Plus Nutrition Shake) from a lick spout within reach of the tongue (Figure 1B). After two days of habituation to the head-fixation and lick spout, we recorded photometry signals from food restricted, hungry mice during self-paced ingestive behavior (Figures 1B and 1C). We observed fluctuations in the activity of CeA^pons^ axons that appeared correlated with bouts of licking and ingestion (Figure 1C). To examine this more closely, we identified these bouts of licking (see Methods) and aligned our recorded photometry signals to the onsets of lick bouts, which revealed reliable increases in activity following the bout onsets (Figures 1D-1F). We then aligned the photometry signals from all lick bouts across all mice to the bout onsets and sorted these by bout duration, which revealed a clear association between lick bout duration and neural activity (Figure 1G). Indeed, across all mice, there was a significant correlation between the photometry signal and lick bout duration, but not with locomotion (Figure 1H). Furthermore, we found that the peak response was similar between bouts of licking with Ensure consumption and dry lick bouts (i.e., without Ensure consumption), supporting a relationship between CeA^pons^ axon activity and the motor action of licking (Figure 1I). The response following lick bout onset was greater in magnitude for longer duration bouts and peaked within 5 seconds, even for lick bout durations lasting much longer than this (Figure 1J). By aligning signals to the end of lick bouts, we also found that responses took several seconds to decrease, which allowed for persistently elevated signals during periods with high likelihood of licking (Figure 1K). When we examined histology of the fiber implants from the mice we recorded, we observed that these photometry signals were similar even when we focused on mice in which the fiber implant passed through the PBN and recorded from the premotor targets below, suggesting that this lick-associated activity is present in CeA axons throughout the dorsolateral pons (Figure S1). Together, these results demonstrate that CeA^pons^ axon activity is highly linked to licking and reflects a state of orofacial motor engagement.

### CeA^pons^ axon activity is independent of hunger state and of palatability, but tail shocks evoke axon response

We asked whether the observed increases in CeA^pons^ axon activity during licking were sensitive to contextual factors including the hunger state of the animal, food palatability, and valence. To determine if CeA^pons^ axon activity reflected hunger state, we compared the photometry signal responses of lick bouts in food restricted and *ad libitum* fed mice during the light cycle (Figures 2A-2C). *Ad lib*. fed mice performed fewer and shorter-duration bouts of licking compared to food restricted mice (Figure 2A), but when we compared lick bouts of similar duration, we found no difference in the photometry response dependent on hunger state (Figures 2B and 2C). Neither the response to the Ensure consumption nor to licking in the absence of Ensure was different between food restricted and *ad lib*. fed mice, further supporting a role for CeA^pons^ axon activity in motor control over state-specific information processing (Figures 2B and 2C).

**Figure 2.**
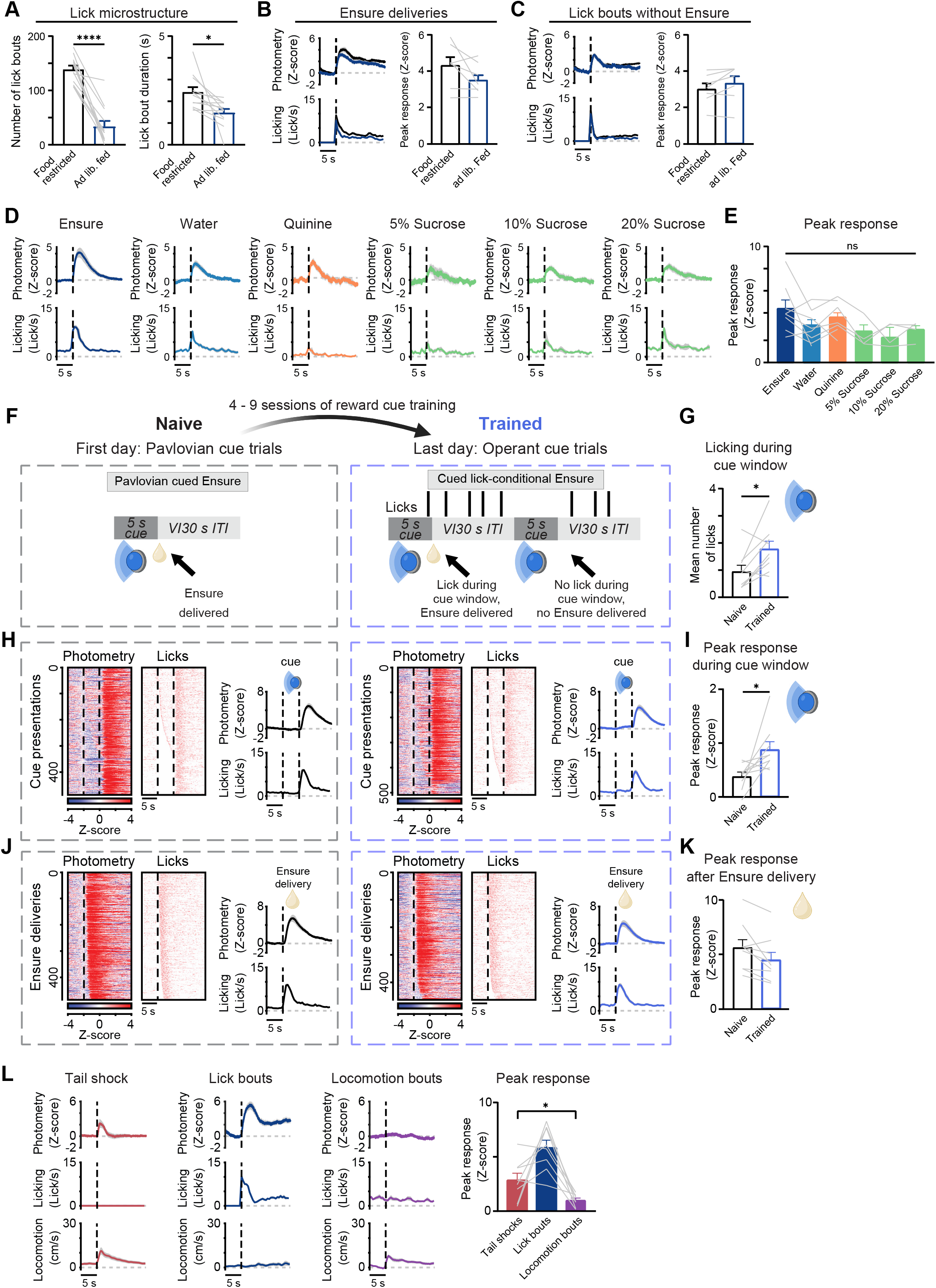
CeA^pons^ axon activity is unaffected by hunger, palatability, or associative conditioning but is activated by aversive stimuli. (A) Left: Mean number of lick bouts during food restricted and *ad lib*. fed sessions. Right: Mean lick bout duration. Two-tailed paired t tests. N = 7 mice (4M, 3F). (B) Left: Time course of mean GCaMP6s photometry response and licking aligned to Ensure deliveries for food restricted and *ad lib*. fed sessions. Right: Peak photometry responses following Ensure delivery. Two-tailed paired t test. N = 7 mice (4M, 3F). (C) Left: Time course of mean photometry and licking aligned to lick bouts lacking Ensure for food restricted and *ad lib*. fed sessions. Right: Peak photometry following lick bout onset. Two-tailed paired t test. N = 7 mice (4M, 3F) (D) Time courses of mean photometry and licking aligned to liquid delivery licking for (in order from left to right): Ensure (N = 8 mice: n = 3M, n = 5F), water (N = 8 mice: 3M, 5F), 1 mM quinine (N = 9 mice: 6M, 3F), 5% sucrose, 10% sucrose, and 20% sucrose (N = 4 mice: 2M, 2F) (E) Peak photometry responses following liquid delivery for the different tastants. Mixed-effects analysis with Dunnett’s correction for multiple comparisons N = 4-9 mice. (F) Left: Trial structure for the first day of Pavlovian conditioning - pairing of a visual cued with Ensure (Naïve). Right: Trial structure for the last day of operant conditioning (Trained). (G) Mean number of licks during cue window on first day of Pavlovian conditioning (Naïve) and last day of operant conditioning (Trained). Two-tailed paired t tests. N = 9 mice (4M, 5F). (H) Heat maps of all trials aligned to cue onset then sorted by latency to first lick during the cue from the Naïve (left) and Trained (right) days. Time courses of photometry and licking averaged across mice aligned to cue onset from the Naïve (left) and Trained (right) days. (I) Peak photometry responses during the cue presentations on first (Naïve) and last (Trained) days of conditioning. Two-tailed paired t tests. N = 9 mice (4M, 5F). (J) Heat maps of all trials aligned to Ensure delivery then sorted by latency to first lick following the delivery from the Naïve (left) and Trained (right) days. Time courses of photometry and licking averaged across mice aligned to Ensure delivery from the Naïve (left) and Trained (right) days. (K) Peak photometry responses after the Ensure delivery presentations on first (Naïve) and last (Trained) days of conditioning. Two-tailed paired t tests. N = 9 mice (4M, 5F). (L) Left: Time courses of mean photometry, licking, & locomotion aligned to tail shocks. Middle, left: Time courses aligned to lick bout onsets for food restricted sessions. Middle, right: Time courses of mean photometry, licking, & locomotion aligned to bouts of locomotion during food restricted sessions. Right: Peak photometry following tail shocks, lick bouts, and locomotion bouts. Repeated-measures one-way ANOVA with Šidák’s correction for multiple comparisons. N = 8 mice (4M, 4F). All data: mean ± SEM.

Next, we assessed if palatability or incentive value of the ingested substance was reflected in the CeA^pons^ axon activity. Altering the liquid delivered from Ensure to either water, bitter quinine, or sucrose of varying concentration did not significantly impact the response (Figures 2D and 2E). As expected, mice licked less upon tasting a high concentration of bitter quinine (1 mM), yet photometry responses remained similar in amplitude to Ensure or water consumption. We suspect that continued reactionary orofacial movements associated with the bitter taste of quinine may be associated with the neural response. Overall, these data add support for linking CeA^pons^ axon activity to the motor actions of licking and ingestion.

CeA neuronal activity is important for aversive and appetitive Pavlovian and operant conditioning^33,34^. Therefore, we asked whether CeA^pons^ axon activity would increase in response to a visual cue that had been associated with Ensure delivery. Mice underwent initial Pavlovian conditioning followed up with operant conditioning in which mice needed to lick during the 5-s cue to receive Ensure at the end of the cue. Following extensive training across 6 to 11 days (150 to 275 total trials), mice licked more during the cue, indicating that they had learned the incentive value of the cue (Figures 2F and 2G). While there was a small, significant increase in the photometry signal during the cue, and this increase aligned well with the onset of licking during the cue (Figures 2H and 2I). Furthermore, we did not find any indication that the activity of these axons was influenced by the prediction or expectation of Ensure delivery, as responses during ingestion of the Ensure remain the same (Figures 2J and 2K). The results are more consistent with CeA^pons^ axon activity being associated with motor aspects of ingestion rather than conditioned sensory stimuli.

Finally, we asked whether an aversive outcome, tail shock, which does not primarily involve orofacial behaviors, but which has previously been shown to drive CeA neuronal activity would lead to increased activity in CeA^pons^ axons^33^. We found that tail shocks drove photometry responses in CeA^pons^ axons, which were smaller in amplitude compared to lick bouts of similar duration to the tail shock-associated locomotion (Figure 2L). Further, since increases in locomotion accompany the tail shock stimulus, we tested whether bouts of locomotion in general lead to increases in CeA^pons^ activity and found that these did not drive strong responses (Figure 2L).

Together, these results suggest that either CeA^pons^ axons display mixed selectivity for licking and tail shocks, or that separate populations of CeA^pons^ axons are selective for these behaviors.

### Distinct CeA^PBN^ axons are active during lick bouts vs tail shocks

To distinguish between these possibilities, we used 2-photon imaging via implanted optical lens to record responses of individual CeA axons and took advantage of the precise z-resolution to restrict our recordings to axons within the PBN only (Figures 3A-3D). We found that, like our photometry results, across all mice we examined, axons and putative synaptic bouton regions of interest (ROIs) were active during bouts of licking and this activity was correlated with the lick bout duration (Figures 3E-3H). We also found ROIs that were responsive during delivery of tail shocks. These ROIs had larger responses following tail shocks compared to ROIs identified based on responses to lick bouts which was consistent across all mice tested (Figure 3I). We compared these lick bout-and shock-preferring ROIs more closely by examining the mean response of each ROI during lick bouts or tail shocks (Figure 3J). This revealed distinct ROIs that were more strongly responsive either to lick bouts or to tail shocks (Figures 3K and 3L). We also found that lick bout-responsive ROIs were more numerous across most fields of view we tested (Figures 3L-3N). Finally, unlike lick bout-preferring ROIs, shock ROIs were more correlated with tail shock-evoked locomotion (Figure 3O). These results extend the photometry findings to the level of individual CeA^PBN^ axons, revealing at least two populations: one preferentially responding to licking and the other to tail shock.

**Figure 3.**
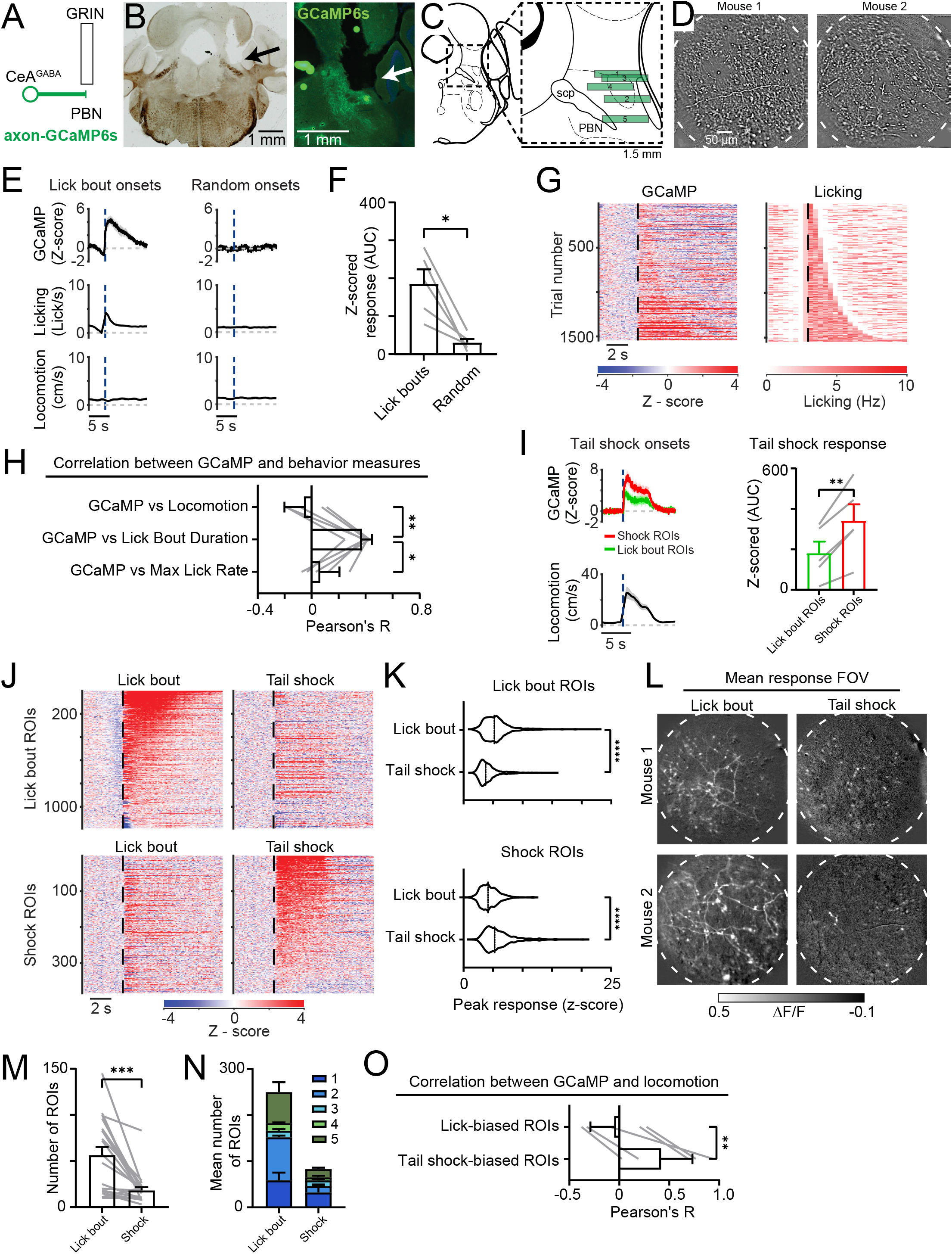
Separate CeA axons within the PBN respond to licking versus tail shocks. (A) Schematic depicting two-photon calcium imaging of CeA axons via an implanted Gradient Refractive Index (GRIN) lens. (B) Left: representative histological brightfield image showing GRIN lens implantation tract (black arrow). Right: epifluorescence image of the GRIN lens implant and calcium sensor expression in CeA axons within PBN (white arrow). (C) Anatomical location within the PBN of the GRIN lens implants of all 5 mice (3M, 2F) included in this dataset. (D) Example fields of view from two mice showing visible CeA axons. Images show the mean baseline-normalized fluorescence signal after local normalization filtering. (E) Left: mean time courses of the calcium signal extracted from lick bout responsive regions of interest (ROIs), licking and locomotion. All signals have been first averaged across all ROIs within each mouse and aligned to lick bout onsets to generate a photometry-lick signal Right: the same dataset but aligned to an equal number of randomized onsets. N = 5 mice. (F) Area under the curve (AUC) Z-scored response following lick bout or random onset. Two-tailed paired t-test. (G) Left: heatmap depicting baseline normalized GCaMP signal (average of all ROIs per mouse) across all possible lick bout onsets sorted by the duration of the lick bouts. Right: heatmap depicting the binned lick rate across all possible lick bouts sorted by the duration of the lick bouts. (H) Pearson’s correlation between GCaMP signal AUC and either locomotion AUC, lick bout duration, or maximum lick rate. Two-tailed paired t-test. N = 5 mice. (I) Left: mean time course of the GCaMP response of lick bout or tail shock preferring ROIs aligned to the onset of tail shocks. Right: AUC of the Z-scored response following lick bout or tail shock onsets. Two-tailed paired t-test. N = 5 mice.

### Optogenetic stimulation of CeA^pons^ axon terminals prolongs ingestive bouts and increases ingestion of aversive tastants

Since CeA^pons^ axon activity correlated with the duration of licking, we asked if stimulating these CeA^pons^ axons would prolong ingestive behaviors. We expressed a red-shift opsin (Chrimson) in GABAergic neurons of the CeA and implanted optic fibers in the dorsal pons, allowing us to optogenetically stimulate axon terminals *in vivo* (Figures 4A and 4B; Figure S2). These mice were trained to reliably consume Ensure from a lick spout. We then photostimulated CeA^pons^ axons terminals only during bouts of licking to test whether stimulation would prolong these bouts (Figure 4C). We found that lick bout-triggered photostimulation increased the duration of lick bouts and total number of licks during a session in a frequency-dependent manner but had no effect on the number of lick bouts (Figure 4D). These results support a role for CeA^pons^ axon activity in maintaining ongoing ingestion. It also suggests that the effects of stimulating CeA^pons^ axons are shorted lived since we did not observe an increase in the number of lick bouts initiated.

**Figure 4.**
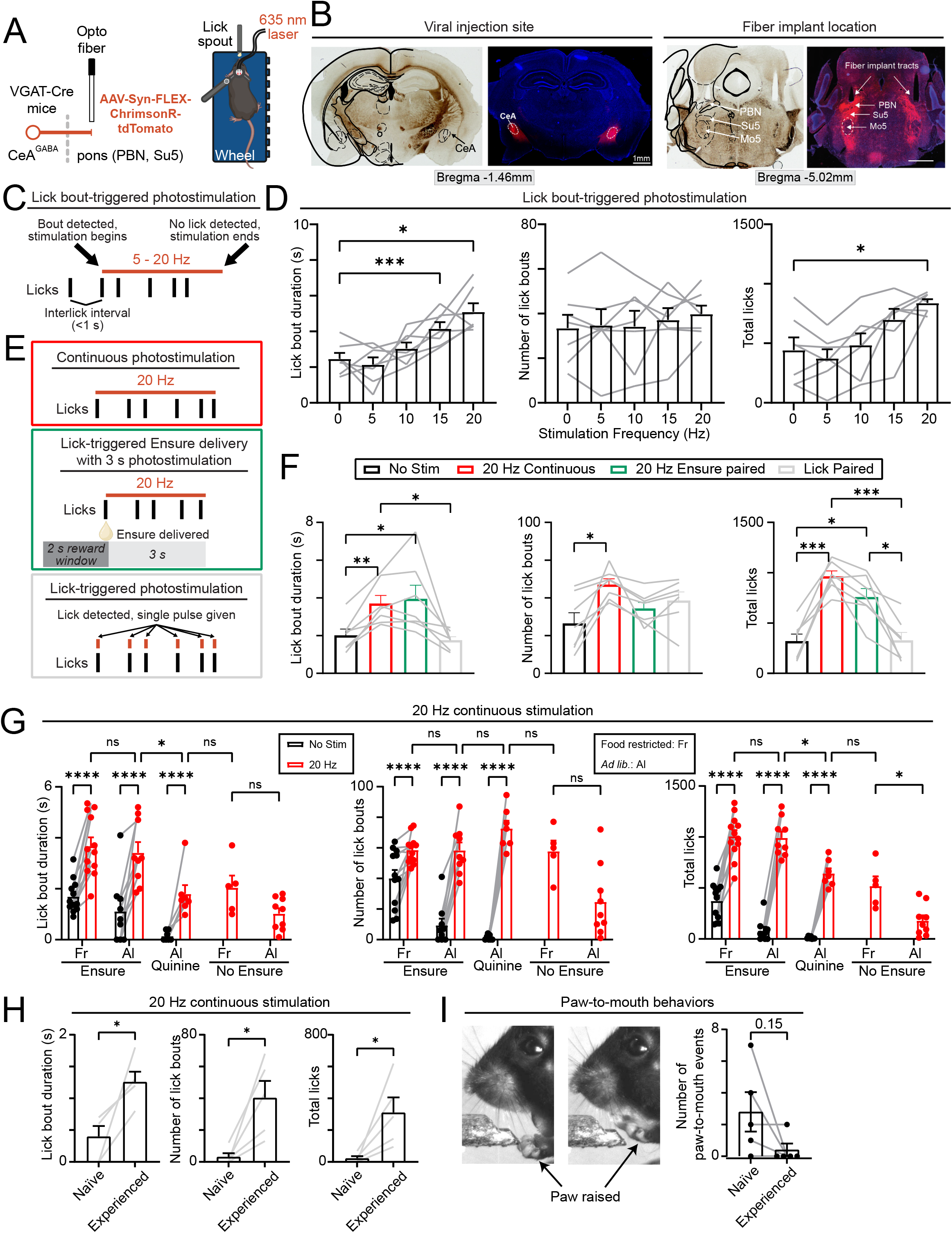
CeA^pons^ photostimulation extends ingestive bouts, drives overconsumption in *ad lib*. fed mice, and overrides aversive taste avoidance. (A) Left: schematic depicting optogenetics experiment in which Cre-dependent ChrimsonR-tdTomato is expressed in GABAergic CeA neurons of VGAT-Cre transgenic mice. Right: schematic of the head-fixed behavioral apparatus and bilateral delivery of red laser light (635 nm). (B) Representative brightfield and epifluorescence images showing ChrimsonR-tdTomato expression at the injection site in the CeA and the axons terminals in the pons below the implanted optogenetics fibers. Note the red-fluorescently label axons in PBN as well as below in the regions such as the supratrigeminal area (Su5) surrounding the trigeminal motor nucleus (Mo5). (C) Schematic describing the lick bout-triggered photostimulation protocol in which pulses (10 ms) of laser light are delivered at 20 Hz following the detection of a lick bout onset and continuing for as long as the interlick interval remains less than 1 s up to a total of 10 s of stimulation. (D) The effect of photostimulation of varying frequencies on lick bout duration (left), number of lick bouts initiated (middle), and total number of licks (right). Repeated-measures one-way ANOVA with Dunnett’s correction for multiple comparisons. N = 7 mice (3M, 4F). (E) Schematics describing continuous 20 Hz photostimulation (red box), 3-s long 20 Hz coincident with the delivery of Ensure (orange box), and single photostimulation pulses (10 ms) paired with each lick detected. (F) The effect of each stimulation protocol compared to no stimulation on lick bout duration (left), number of lick bouts initiated (middle), and total number of licks (right). Repeated-measures one-way ANOVA with Tukey’s correction for multiple comparisons. N = 7 mice (3M, 4F). (G) The effect of 20 Hz continuous stimulation on lick bout duration (left), number of lick bouts initiated (middle), and total number of licks (right). This effect was test when mice were food restricted (Fr) or *ad lib*. fed. (Al) and given either Ensure, quinine (1 mM), or nothing (No Ensure) to ingest. Two-way ANOVA with Šidák’s correction for multiple comparisons. N = 12 (6M, 6F) food restricted mice offered ensure with no stim or stim. N = 9 (5M, 4F) *ad lib*. fed mice offered ensure with no stim or stim. N = 7 (3M, 4F) *ad lib*. fed mice offered quinine with no stim or stim. N = 5 (3M, 2F) food restricted mice offered nothing during stim. N = 9 (5M, 4F) *ad lib*. fed mice offered nothing during stim. (H) The effect of 20 Hz continuous stimulation on lick bout duration (left), number of lick bouts initiated (middle), and total number of licks (right) in mice exposed to a lick spout for the first time (Naïve) versus mice Experienced with licking at a lick spout to receive Ensure. The lick spout did not contain any Ensure for these experiments and mice were *ad lib*. fed. Two-tailed paired t-test. N = 5 mice (3M, 2F). (I) Left: images depicting paw raises to the mouth that were observed during continuous 20 Hz photostimulation. Right: quantification of the number of paw-to-mouth events detected in Naïve versus Experienced mice. Two-tailed paired t-test. N = 5 mice (3M, 2F).

Could stimulating CeA^pons^ axon activity drive the initiation of ingestion? To address this question, we tested whether continuously stimulating CeA^pons^ axons would increase the number of lick bouts (Figure 4E). We found that continuous 20 Hz stimulation caused an increase in the number of lick bouts, lick bout duration, and the total number of licks (Figure 4F). In contrast, stimulating CeA^pons^ axons for 3 seconds just following a delivery of Ensure increased lick bout duration and total licks without influencing the number of lick bouts, similar to our findings when we stimulated in closed-loop coincident with bouts of licking (Figures 4E and 4F). Finally, we asked whether stimulating CeA^pons^ axons paired with each individual lick would influence this behavior (Figure 4E). We found that there was no change in any parameter, demonstrating that one-to-one stimulation with ongoing licking had no effect on future behavior (Figure 4F). Together, these results show that stimulating CeA^pons^ axons increases the likelihood of licking, the ability to sustain ongoing bouts of licking, and that these effects are short lasting.

We next investigated whether factors like satiety, taste, ingestion, and training altered the effect of stimulating CeA^pons^ axons. We tested whether *ad lib*. fed mice, which are less motivated to lick for ensure, would be resistant to CeA^pons^ axon stimulation and found that both bilateral and unilateral photostimulation drove licking behavior (Figure 4G and Figure S3A). We tested a bitter tastant, quinine (1 mM), which mice strongly avoid consuming and found that stimulating CeA^pons^ axons was able to induce licking and consumption of quinine, but not to the level of Ensure (Figure 4G). Even though photostimulation caused a similar number of lick bouts to be initiated, lick bout durations were shorter with quinine compared to Ensure, consistent with the taste being unpalatable and indicating that taste can still influence licking behavior during CeA^pons^ photostimulation. Finally, we asked how removing all liquid delivery while maintaining the tactile sensation of the lick spout would impact licking behavior. We found that CeA^pons^ photostimulation was less effective at driving prolonged lick bouts, like the results with quinine, indicating that ingestion of the palatable Ensure was necessary to maintain long lasting lick bouts (Figure 4G).

Finally, we compared photostimulation in *ad lib*. fed mice that were only habituated to head-fixation but not to a lick spout (Naïve) to mice that were well experienced at licking for Ensure (Experienced). We found that Naïve mice were less likely to display licking behavior and more likely to display putative eating-associated behaviors and other orofacial movements compared to Experienced mice (Figures 4H and 4I). Together, these results indicate that CeA^pons^ axons stimulation influences orofacial movements, prolongs lick bouts, and causes overconsumption in *ad lib*. fed mice, but that licking behavior still depends on converging information including taste, ingestion, and training.

### CeA^pons^ axon stimulation induces orofacial movements and motor programs of ingestion

We wanted to further understand the orofacial movements we observed in Naïve mice, which suggested that the exact orofacial behaviors performed may depend on specific contextual information. To explore this, we used videography, pose estimation (DeepLabCut), and classification (SimBA) of orofacial behaviors (Figures 5A and 5B). In the absence of any lick spout or food, photostimulation of CeA^pons^ axons increased the number of tongue protrusions into the air in a frequency-dependent manner (Figures 5C and 5D; Figure S3B; Movie S1). We also observed movements of the jaw during 20 Hz photostimulation that appeared rhythmic (Figure 5E; Movie S1). To analyze these jaw movements, we excluded frames with licks and found that 20 Hz photostimulation increased the total distance the jaw moved (Figure 5F; Figure S3C; Movie S1). Together, these results demonstrate that photostimulating CeA^pons^ axons causes overt orofacial movements in the absence of ingestion.

**Figure 5.**
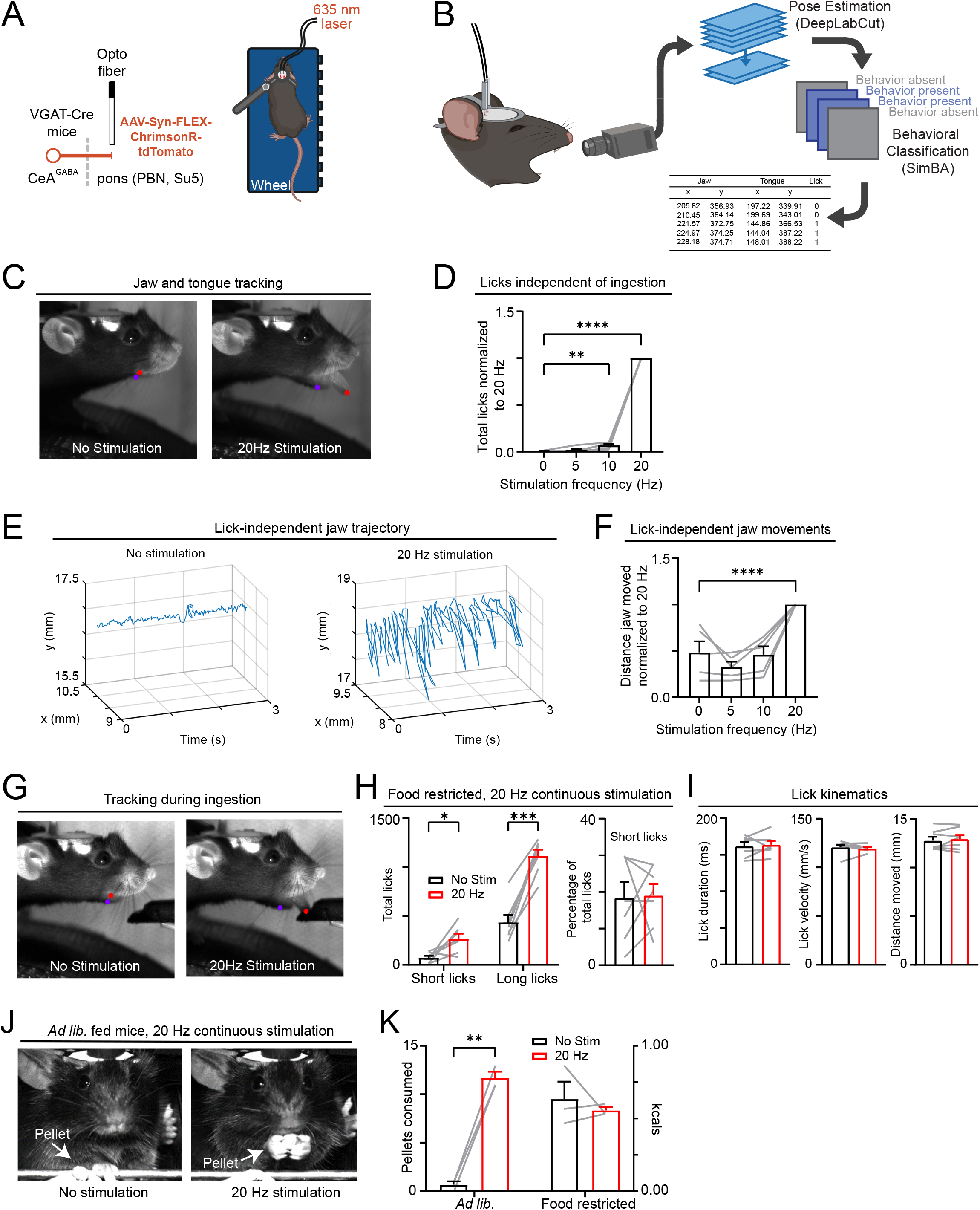
Photostimulation of CeA^pons^ axons promotes orofacial ingestive behaviors without altering behavior characteristics. (A) Left: schematic depicting optogenetics experiment in which Cre-dependent ChrimsonR-tdTomato is expressed in GABAergic CeA neurons of VGAT-Cre transgenic mice. Right: schematic of the head-fixed behavioral apparatus and bilateral delivery of red laser light (635 nm; 5 mW). (B) Schematic of workflow for capturing and quantifying orofacial behaviors from videography to pose estimation (DeepLabCut) to classification (SimBA). (C) Representative images depicting key point tracking of lower jaw (purple dot) and the tip of the tongue (red dot) from videography of mice in the absence of any lick spout. Images highlight orofacial movement and tongue extension observed with 20 Hz continuous photostimulation. (D) Analysis of licks detected by videography in *ad lib*. fed mice during photostimulation of varying frequency. Results are normalized within mouse to the total licks detected with 20 Hz stimulation. Repeated-measures one-way ANOVA with Dunnett’s correction for multiple comparisons. N = 5 mice (3M, 2F). (E) Representative trajectory of the lower jaw with or without 20 Hz photostimulation. Jaw movement related to licking has been removed. (F) Analysis of jaw movement distance measured by videography in *ad lib*. fed mice during photostimulation of varying frequency. Results are normalized within mouse to the total distance measured with 20 Hz stimulation. Jaw movement related to tongue extension is not included in this analysis. Repeated-measures one-way ANOVA with Dunnett’s correction for multiple comparisons. N = 5 mice (3M, 2F). (G) Representative images depicting key point tracking of lower jaw (purple dot) and the tip of the tongue (red dot) from videography of mice engaged in licking to consume Ensure. Images highlight orofacial movement and tongue extension observed with 20 Hz continuous photostimulation. (H) The effect of 20 Hz photostimulation on total licks of food restricted mice analyzed from videography. Left: both short licks that do not reach the lickspout and long licks that contact the lick spout are increased. Repeated-measures two-way ANOVA with Šidák’s correction for multiple comparisons. N = 7 mice (3M, 4F). Right: the percentage of short licks out of all licks did not change with photostimulation. Two-tailed paired t-test. (I) Analysis of lick movement characteristics. Two-tailed paired t-test. N = 7 mice (3M, 4F). (J) Representative images from videography of head-fixed pellet grasping behavior. Images highlight pellet grasping and consumption by an *ad lib*. fed mouse observed with 20 Hz continuous photostimulation. (K) The effect of 20 Hz photostimulation on total pellets consumption by *ad lib*. fed and food restricted mice. Two-way ANOVA with Šidák’s correction for multiple comparisons. N = 3 mice.

To examine how stimulating CeA^pons^ axons influenced the motor characteristics of ongoing natural licking behavior, we analyzed the tongue protrusions of mice that reliably licked at a lick spout to ingest Ensure (Figure 5G). We found that photostimulation of CeA^pons^ axons increased the total number of licks, including previously undetected short licks that did not contact the lick spout, but the proportion of short to long licks did not change, indicating a general increase in all lick types (Figure 5H). Furthermore, we did not find a difference in the velocity or total distance of the tongue extension during stimulation versus no stimulation sessions (Figure 5I). These data indicate that stimulating CeA^pons^ axons increases the likelihood of tongue extensions in mice experienced with licking to consume Ensure without a detectable impact on the motor characteristics of the behavior.

We also noted other putative eating-associated behaviors including paw raises to the mouth, especially in mice naïve to licking at the lick spout for Ensure (Figure 4I). To examine these types of ingestive behaviors in head-fixed mice, we modified our apparatus to allow mice to grasp small chow pellets and eat solid food^35^ (Figure 5J). We found that in *ad lib*. fed mice that were trained to grasp and eat pellets, photostimulating CeA^pons^ axons led to the overconsumption of pellets (Figure 5J and 5K; Movie S2). Together, these results support a role for CeA^pons^ axons in promoting the likelihood of ingestive motor programs, without specifying the exact motor patterns or actions.

### CeA^pons^ axon photostimulation increases consumption of nearby food and non-food objects even at the cost of potential starvation

To extend our findings that CeA^pons^ axon activation increases orofacial behaviors and ingestion without dictating specific motor actions, we tested ingestive behaviors in freely moving mice (Figure 6A and Figure S4). We first examined *ad lib*. fed mice, which we expected would show increased eating during photostimulation. Surprisingly, we observed a decrease in the food consumed during continuous optogenetic stimulation compared to the same period in sessions without stimulation (Figure 6B). Having observed that CeA^pons^ axon stimulation paradoxically suppressed food consumption, we next tested whether fasted mice, which typically consume a large amount food, would demonstrate similarly suppressed consumption. We found that fasted animals ate almost none of the available chow during 20 Hz continuous photostimulation, and this effect depended on the stimulation frequency (Figure 6C and Figure S4A). Control mice, in which Chrimson was not delivered to CeA neurons, did not show suppressed eating during photostimulation (Figure S4B). Upon further examination of the videography, we observed that mice were actively engaged in ingestive behaviors during photostimulation but appeared to be consuming the bedding rather than approaching the available food (Movie S3). These results suggest that photostimulation timing and proximity to actual food may be critical and that ingestive motor behaviors triggered by stimulating CeA^pons^ axons in freely moving mice is indiscriminate to whether objects are truly food.

**Figure 6.**
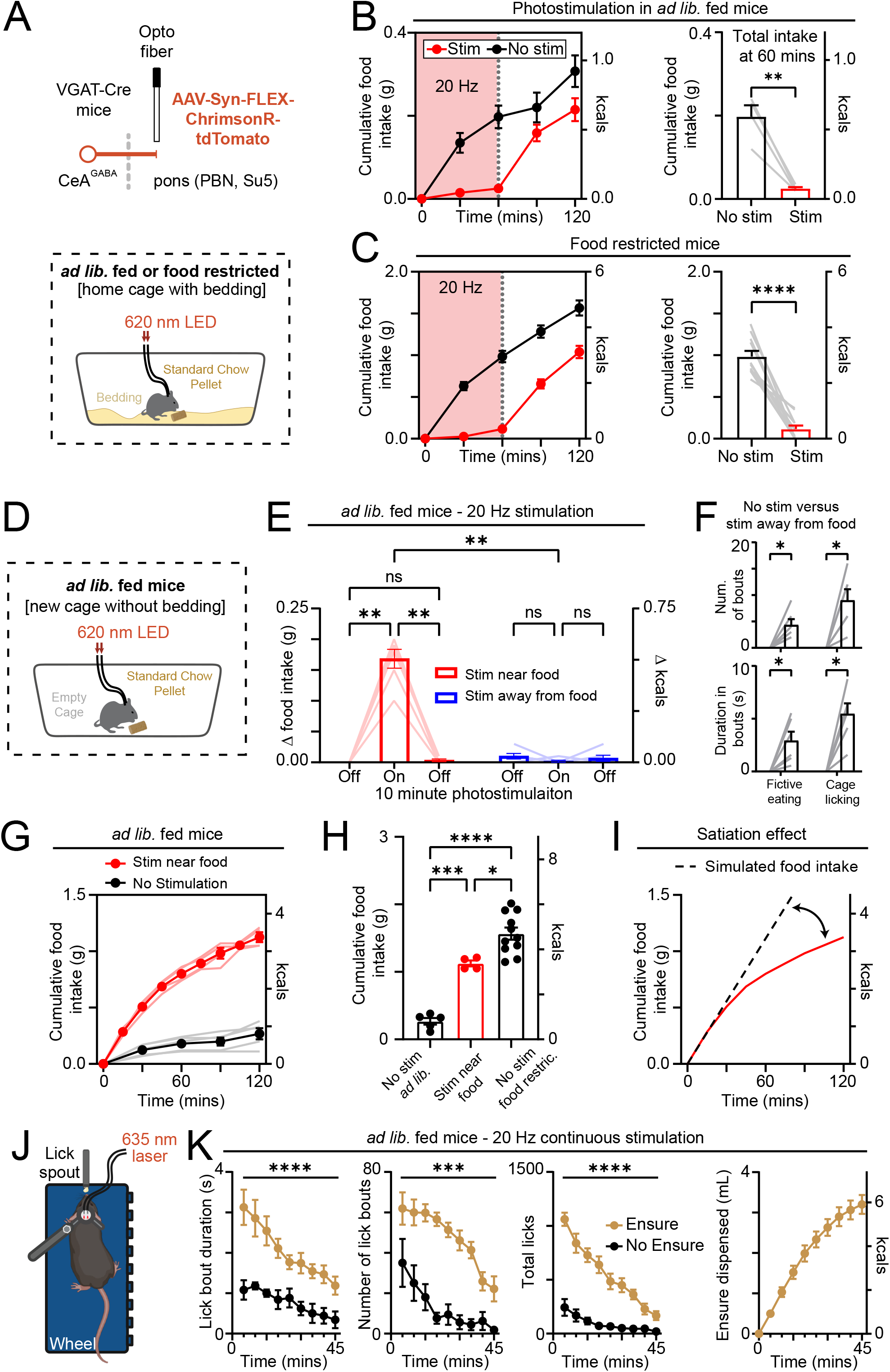
CeA^pons^ photostimulation promotes ingestive behaviors directed at nearby objects. (A) Top: schematic depicting optogenetics experiment in which Cre-dependent ChrimsonR-tdTomato is expressed in GABAergic CeA neurons of VGAT-Cre transgenic mice. Bottom: schematic of the freely moving, home cage feeding assay with bilateral delivery of red LED light (620 nm). (B) Left: average time course of food intake of *ad lib*. fed mice over 2 hours with either 20 Hz continuous or no photostimulation during the first 60 minutes. Right: total food consumed in the first 60 mins with or without photostimulation. Two-tailed paired t-test. N = 4 mice (1M, 3F). (C) Left: time course of average food intake of food restricted mice over 2 hours with either 20 Hz continuous or no photostimulation during the first 60 minutes. Right: total food consumed in the first 60 minutes with or without photostimulation. Two-tailed paired t-test. N = 11 mice (5M, 6F). (D) Schematic of the freely moving, empty cage feeding assay with bilateral delivery of red LED light (620 nm). (E) Food consumed during periods with or without 20 Hz photostimulation either when the mouse was near the food (red) or away from the food (blue). Repeated-measures two-way ANOVA with Šidák’s correction for multiple comparisons. N = 6 mice (4M, 2F). (F) Top: number of bouts of fictive eating or cage licking detected with or without photostimulation while the mouse is away from the food. Bottom: duration in bouts of fictive eating or cage licking detected with or without photostimulation while the mouse is away from the food. Repeated-measures two-way ANOVA with Šidák’s correction for multiple comparisons. N = 6 mice (4M, 2F). (G) Time course of average food intake of *ad lib*. fed mice over 2 hours with either 20 Hz stimulation (N = 4 mice; 2M, 2F) near food or no photostimulation (N = 5 mice; 5F). (H) Food intake of *ad lib*. fed mice receiving no stimulation (N = 5) or 20 Hz stimulation when near food (N = 4), and food restricted mice (N = 11) receiving no photostimulation. Two-way ANOVA with Tukey’s correction for multiple comparisons. (I) Representation of food intake simulated to increase linearly during 2 hours of photostimulation compared to results obtained with 20 Hz photostimulation when mice are nearby food. Note the predicted effect of satiation signals slowing down the rate of consumption. (J) Schematic of the head-fixed behavioral apparatus with available liquid food (Ensure) and bilateral delivery of red laser light (635 nm). (K) The effect of continuous 20 Hz photostimulation on lick bout duration (left), number of lick bouts initiated (middle, left), total number of licks (middle, right), and Ensure dispensed (right) across 45 minutes. Mixed-effects model. n = 9 mice (5M, 4F) with Ensure, n = 5 mice (3M, 2F) without Ensure.

Indeed, when we photostimulated CeA^pons^ axons in water restricted mice that were given a small, plastic petri dish containing water, we similarly observed a decrease in ingestion of the needed water and an increase in licking behaviors directed to the walls of the cage (Figure S4C). We also observed chewing and ingestion of the plastic petri dish consistent with our prior observations that photostimulation drives ingestion of nearby non-food objects (Figure S4D). To continue exploring these non-food-directed orofacial behaviors, we tested whether photostimulating CeA^pons^ axons would affect behaviors directed at a moving toy bug, which has been used as an artificial prey when investigating the role of CeA in predatory hunting^16^. We found that mice actively avoided the toy bug in the absence of photostimulation, but upon photostimulation of CeA^pons^ axons, mice would bite and remain clenched on the prey if it came near them (Figures S4E and S4F; Movie S4). Together these results show that stimulating CeA^pons^ axons commands ingestive orofacial behaviors that are directed towards proximate objects and the specific motor action—licking, chewing, or biting—seems to be dictated by the locally available sensory information.

To test this hypothesis, we designed a closed-loop assay with stimulation occurring either while the mouse was nearby a food pellet or on separate sessions while the mouse was away from food (Figure 6D). The experiment was conducted in a cage without any bedding to reduce ingestion of non-food objects. As predicted, we found that if photostimulation was timed to occur when the mouse approached and was nearby food, it induced robust chewing and ingestion of the pellet (Figure 6E; Movie S5 and S6). In contrast, when photostimulation occurred while the mouse was away from the pellet, it resulted in chewing and licking behaviors directed at the cage floor or the air (i.e., raising paws to mouth to fictively eat a non-existent pellet) and did not result in consumption of the pellet (Figures 6E and 6F; Movie S7). These putative eating-associated behaviors were accompanied by a decrease in locomotion consistent with this stimulation favoring a behavioral state of exploitation of local food over foraging for new food (Figure S4G).

### Cessation of consumption remains intact despite CeA^pons^ axon optogenetic stimulation

Having discovered that the timing of the photostimulation played a critical role, we next asked whether recurrently photostimulating CeA^Pons^ axons when the mouse was near a food pellet could overcome satiety and drive hours-long overconsumption in *ad lib*. fed mice. We found that while this closed-loop stimulation paradigm did drive robust overconsumption almost to the level of a food restricted mouse, the rate of consumption tapered off over the course of two hours (Figures 6G and 6H; Movie S5). Indeed, we observed that as the experiment continued, mice spent less time approaching the pellet, while overall distance moved did not change, consistent with a switch from food seeking to active food avoidance (Figure S4H). We also tested if this avoidance of consumption with prolonged ingestion would occur even in the head-fixed context where mice could not choose to physically distance themselves from the ingestive substance. Indeed, we found that head-fixed mice decreased licking, and the amount of Ensure consumed over the course of 45 minutes of continuous stimulation was consistent with visceral satiation signals arresting the behavior (Figure 6J and 6K). However, when we repeated this experiment without any Ensure available for consumption, we also observed a decrease in licking behavior, indicating that other factors like fatigue may also contribute.

Finally, we tested whether mice would prefer or avoid CeA^pons^ photostimulation. Prior studies have found that activating certain populations of CeA neurons can drive an immediate place preference^12^. Using a real-time placed preference (RTPP) assay, we found that, consistent with a preference for CeA^pons^ photostimulation, mice spent more time on the side of the chamber that led to photostimulation on all three days of testing (Figure S5A and S5B). We did not observe any preference for the side of the chamber paired with photostimulation in the absence of stimulation on the third day, suggesting that two days of conditioning did not lead a learned preference for photostimulating CeA^pons^ axons (Figure S5B). However, upon examination of the behavioral videography, we found that mice were engaged in active chewing behaviors directed at the walls and the plastic divider of the RTPP chamber, which also resulted in a decrease in the velocity of the mice (Figures S5C and S5D). This decreased velocity and orofacial behavior is consistent with our results during stimulation in the home cage (Figure 6F and Figure S4G). Taking all these results into consideration, the findings suggest that while mice do not actively avoid CeA^pons^ axon photostimulation, they become less mobile and are preoccupied with consummatory behaviors (i.e. chewing and licking) that sustain the photostimulation, making it difficult to interpret this stimulation as being actively preferred. Taken together with our freely-moving feeding assays, these RTPP results suggest that activation of CeA^pons^ axons drives ingestive behaviors toward the nearest accessible items—be they food or plastic.

## DISCUSSION

We originally set out to test the hypothesis that CeA projections to the parabrachial nucleus (CeA^PBN^) promote feeding by suppressing satiety signals or enhancing food palatability, as suggested by previous studies^12^. We were surprised to find extensive CeA terminals within adjacent premotor nuclei as these are typically not displayed and discussed in studies investigating this pathway. These broad CeA projections made it challenging to specifically attribute results to a specific pontine target, although we used two photon imaging and post hoc histological assessments to gain some anatomical insights. Overall, our results suggest that activation of CeA^pons^ axons promotes feeding behavior likely by modulating the orofacial motor system, which occurred even in the absence of food or metabolic need. While CeA outputs have been implicated in suppressing PBN-mediated pain signals^36^, we found that stimulation triggers rhythmic licking, biting, and chewing. It also prioritizes these consummatory behaviors over food seeking behaviors, which would not be expected from just suppressing satiety. Instead, we propose that CeA axons directly engage premotor circuits adjacent to the PBN, which disinhibits the orofacial motor circuits to promote consumption.

By combining fiber photometry and two-photon imaging, we show that CeA^pons^ axon activity scales with ongoing ingestive motor action. Lick-related signals grew stronger with longer bout durations and were observed even in the absence of food or liquid, including during licking of dry spouts. These activity patterns were largely insensitive to hunger state or palatability, suggesting that CeA^pons^ output reflects motor engagement rather than food value or internal drive. Two-photon imaging of signals confirmed to be from axons within the PBN reproduced our photometry results. It also revealed that lick-related and aversion-related responses are found in separate axons, supporting the existence of functionally specialized subpopulations within this projection-defined pathway. However, lick-preferring axons were more numerous than aversion-related axons, indicating why activation of all axons would lead to an overall increase in feeding behaviors.

Optogenetic stimulation of CeA^PBN^ axons confirmed a causal role for this pathway in driving ingestion. Activation during ongoing feeding bouts extended consumption, and, in both freely moving and head-fixed animals, stimulation triggered rhythmic orofacial actions such as licking, biting, and chewing. These behaviors have been noted in prior studies^12,14,15,22^, although they have not been closely investigated and the possibility that these behaviors are mediated by pontine orofacial premotor neurons has not been discussed. These behaviors occurred even in the absence of food and were frequently directed toward nearby inedible objects, indicating that the stimulation was sufficient to activate motor programs associated with feeding, independent of sensory input or motivational context. In some cases, stimulation caused mice to prioritize ingesting non-food stimuli over accessible food leading to a paradoxical decrease in food consumption in hungry mice. Interestingly, this decrease in food intake with CeA neuron activation has been reported previously^15^ and our results, especially our closed-loop photostimulation experiments, provide a mechanism by which mice prioritize local ingestive behaviors over goal-directed food seeking. This finding suggests that CeA^pons^ activation may impair the ability to appropriately target ingestive behavior, instead favoring orofacial behaviors guided by only very proximal cues and is unlikely to be causing this effect by suppressing satiety signals.

### CeA^pons^ activation promotes consummatory motor programs rather than specific motor actions

We attempted to distinguished whether CeA^pons^ activation impacted the specific motor patterns or instead more generally created a state in which any ingestive behavior was more likely to occur. Our analysis of the specific kinematics of licking behavior suggested that stimulation did not on average alter normal motor action characteristics such as the velocity of the movement or the length of the tongue extension, but did increase the likelihood of context-guided orofacial movements including licking for Ensure or reaching to grasp food pellets. Further supporting this, mice tended to perform a variety of orofacial behaviors including putative eating-associated behaviors like paw-to-mouth movements during CeA^pons^ stimulation if mice were not previously habituated to the lick spout (i.e., naïve). However, upon training to lick for Ensure at the lick spout, the experienced mice selectively increased licking during CeA^pons^ stimulation, whereas mice trained to eat food pellets selectively increased pellet grasping, but not licking behaviors. These results are consistent with a role for CeA^pons^ in disinhibiting orofacial motor circuits, possibly by inhibiting the supratrigeminal neurons that tonically inhibit motor neurons^31^, instead of commanding specific motor actions.

### Which pontine circuits likely mediate these motor actions?

The PBN is situated near the pontine supratrigeminal and peritrigeminal premotor nuclei^37^, which surround the trigeminal motor nucleus just ventral to the PBN. These regions contain putative central pattern generator neural circuits that coordinate the orofacial muscle groups^30,37^. In addition, inhibitory neurons of the supratrigeminal nucleus directly synapse onto and maintain a tonic inhibition of trigeminal motor neurons^31^. While CeA projections to the medulla have previously been linked to direct jaw muscle engagement and to predatory hunting behaviors without affecting feeding^16^, our results show that stimulation of CeA^pons^ axons can elicit similar motor actions in a feeding context, leading to overconsumption of food and ingestion of inedible objects. Thus, we believe the most parsimonious interpretation of our results is that CeA^pons^ stimulation inhibits supratrigeminal neurons that maintain the tonic inhibition of orofacial motor neurons.

Interestingly, we observed an array of motor programs were able to be driven by the CeA^pons^ pathway including licking and arm movements, which are controlled by motor nuclei not located in the pons^38,39^. We also observed biting and capturing of artificial prey similar to hunting behaviors observed when stimulating CeA projections to the medulla^16^. We suspect that either CeA neurons send collateral axons throughout the reticular formation or that polysynaptic connections between pontine and medullary reticular formation circuits may allow for a wider array of ingestive behaviors to be facilitated. Taken together, our findings refine the interpretation of CeA^pons^ function by highlighting a motor facilitation component that likely contributes to the robust overconsumption effects reported in prior work.

### What might be the purpose of the motor-associated signals in CeA^PBN^ pathway?

We hypothesize that CeA may concurrently influence both visceral sensory processing and orofacial motor control. This might serve the purpose of either a motor efference copy^40^ that is relayed to the viscerosensory PBN pathway to negate already known sensory information, or alternatively to act as a mechanism to efficiently sustain behavior by ignoring competing signals relayed via the PBN such pain or nausea related to stomach distension. We hypothesize that selective manipulations of CeA terminals solely onto PBN neurons may prolong meals or increase food seeking behavior but should not initiate consummatory motor programs. While it is possible that CeA neurons may inhibit PBN neurons that project to orofacial motor circuits, such a pathway has not been observed. Future studies using simultaneous recording of CeA axons and PBN neurons may help distinguish the role of these collaterals.

### CeA^pons^ stimulation does not override homeostatic constraints indefinitely

CeA^pons^ stimulation drives ingestive behaviors, but these behaviors eventually subside, suggesting they remain sensitive to internal state. In freely moving *ad lib*. fed mice, closed-loop stimulation of CeA^pons^ axons initially triggered robust consummatory responses, but after substantial food intake, mice began avoiding the pellet, thereby preventing further stimulation. We speculated that this avoidance reflects visceral satiety signals, such as gastric distension, that oppose continued consumption. To test whether this effect depended on the ability to avoid food, we repeated the experiment in head-fixed mice consuming liquid Ensure in which the lickspout was always within reach of the tongue. Even in this condition, mice stopped licking in response to stimulation over time, demonstrating that CeA^pons^-driven orofacial behaviors can be suppressed. Notably, when stimulation was delivered without any food present, licking still decreased over time, suggesting the effect was not solely due to gastric distension. This decline may reflect response fatigue, synaptic adaptation, or a learned suppression of behavior. Future work should dissect the circuit mechanisms that limit persistent CeA^pons^-driven ingestion, which may reveal insights into the circuits capable of controlling ingestion.

### Broader implications

These findings also broaden our understanding of how emotional and motivational states can influence feeding by acting not only on circuits that regulate food value or internal need, but also on the motor systems that generate ingestive behavior. Rather than solely enhancing the salience of food or suppressing aversive input, CeA projections are capable of directly releasing patterned orofacial actions associated with consumption. This perspective helps explain how feeding can occur in the absence of hunger, and why emotionally driven states—such as stress or anxiety— may lead to maladaptive overconsumption. The observation that mice engaged in ingestive behaviors toward inedible objects, even when food was available, suggests a breakdown in sensory-motor coordination that mirrors aspects of compulsive or pica-like behavior. These behaviors may extend beyond just feeding to other orofacial behaviors such as nail biting or teeth grinding (bruxism), which are often triggered or exacerbated by emotional states^41^. In this view, CeA outputs may serve as a general pathway through which affective signals translate into patterned motor actions, sometimes with maladaptive outcomes.

### Limitations of the study and future directions

While our findings reveal a clear role for CeA projections in promoting orofacial motor behaviors, several questions remain about the specificity and downstream organization of this pathway. Our optogenetic approach likely co-activated axons collateralizing to both sensory and motor-related targets, making it difficult to fully disentangle the relative contributions of satiety suppression versus motor disinhibition. Future studies will need to determine the amount of collateralization between these brainstem targets and attempt to selectively target them. Moreover, although our imaging data revealed functional segregation of lick-and aversion-related signals within distinct axons, future work using intersectional or activity-tagging approaches will be needed to map specific cell types to their behavioral effects^14^. We also found that hunger state influenced the behavioral impact of CeA^pons^ stimulation— suggesting that internal state can gate the expression of these motor patterns—but we did not directly test how emotional or stress-related states might modulate this circuit’s function. Additionally, we were unable to establish whether and when this pathway is necessary for feeding behavior, although other studies have provided a causal role of the CeA during eating, drinking, and hunting^12,15,16,42^. Given its apparent role in overriding internal cues and driving behavior toward inappropriate targets, it is likely that CeA^pons^ projections become more relevant in contexts in which the affective state influences ingestion, rather than during homeostatically driven feeding. Finally, our results highlight a broader challenge in interpreting functional manipulations of projection-defined circuits: even when targeting anatomically specific outputs like CeA^PBN^, collateralization can lead to behavioral effects that reflect broader network engagement than intended.

### Conclusions

Together, our findings reveal that CeA projections to the pons promote feeding behavior not only by modulating sensory feedback, but by engaging motor circuits that drive ingestion. This shifts the interpretation of CeA^pons^ function from a purely modulatory role in satiety and aversion toward a more active role in facilitating orofacial action patterns. By demonstrating that emotional circuits can disinhibit specific motor programs, our study highlights a mechanism through which affective states may override homeostatic control to drive maladaptive consumption. More broadly, this work underscores the importance of considering motor circuit engagement in limbic control of behavior, offering new insight into how emotional states can shape patterned actions across different behavioral domains.

## SUPPLEMENTAL INFORMATION

## RESOURCE AVAILABILITY

### Lead contact

Further information and requests for resources and reagents should be directed to and will be fulfilled by the lead contact, Andrew Lutas.

### Materials availability

This study did not generate any new materials.

### Data and code availability

All original data and code will be deposited and publicly available after publication. Any additional information required to reanalyze the data reported in this paper is available from the lead contact upon request.

## ACKNOWLEDGMENTS

This work was supported by the National Institute of Diabetes and Digestive and Kidney Disorders (NIDDK) and Division of Veterinarian Research (DVR) animal facility staff. We thank J. Barrett for advice with head-fixed grasping studies. We thank A. Levine and M. Reitman for helpful discussions and feedback. We also thank M. Krashes, H. Tejeda, Y. Carrasquillo, M. Penzo, N. Ryba, S. Zhang, and M. Andermann for providing suggestions. We are grateful to members of the Krashes, Reitman, and Lutas labs for support. This research was supported by the Intramural Research Program of the NIH, The National Institute of Diabetes and Digestive and Kidney Diseases (1ZIA-DK075168).

## AUTHOR CONTRIBUTIONS

Conceptualization: D.L., A.L; Methodology: D.L., J.W., and A.L; Investigation: D.L., J.I., J.W., A.P., L.R., and A.L.; Software: A.L.; Validation: D.L., J.I., J.W. and A.L.; Formal Analysis and Data Visualization: D.L., J.I., J.W., and A.L.; Resources: A.L.; Writing—original draft: D.L. and A.L.; Writing—review & editing: D.L., J.W., and A.L., Funding Acquisition: A.L.; Supervision: A.L.

## DECLARATION OF INTERESTS

The authors have no competing interests to declare.

Supplemental Figures S1-5 Supplemental Movies S1-7

## EXPERIMENTAL MODEL AND SUBJECT DETAILS

All animal care and experimental procedures were approved by the National Institutes of Health Animal Care and Use Committee. All experiments were performed on adult Vgat-ires-cre mice bred and maintained in our mouse colony (original Jackson Lab Strain # 028862). Mice were singly housed in 22-24°C with a 12-hour light/12-hour dark cycle. Mice were fed with *ad libitum* standard chow (Teklad F6 Rodent Diet 8664) and given *ad lib*. access to water unless otherwise stated for experimental purposes. For experiments involving food restriction, mice received ~3 g of standard chow per day, and for experiments implementing water restriction, mice were given access to water for a minimum of 1 h each day. In both cases, body weight was monitored daily to maintain body weight at 85-90% of free-feeding weight. Mice used for surgery were at least 8 weeks old at the time of surgery. Mice of both sexes were used for all experiments. Details on the number of mice used per experiment are found in the figure legends.

## METHOD DETAILS

### Stereotaxic surgery and viral injections

Surgeries were performed as previously described^1,2^. Briefly, mice were anesthetized with 3-4% isoflurane first in an induction chamber then mounted on a stereotaxic apparatus (Kopf, Model 940). Mice were placed on a heating pad (Stoelting, Rodent Warmer X1) set at 37°C then subcutaneously injected with 4 mg/kg meloxicam slow release and 0.5-1.0 mL 0.9% sterile saline. The skull was leveled, and craniotomy was performed. For viral injections into the CeA, the following coordinates relative to Bregma were used: AP −1.2 mm, ML ± 2.6 mm, DV −4.9 mm. Viruses were injected at a rate of <50 nL/min using a glass pipette with a tip diameter of 20-40 μm attached to an air pressure system, and the pipette was withdrawn 5-10 min after injection to allow for diffusion of virus. Optic fibers or GRIN lens implantations were secured to the skull using C&B Metabond (Parkell), along with a custom-made titanium headpost (H.E. Parmer)^3^. A minimum of 3 weeks elapsed post-surgery to allow mice to recover and to allow for sufficient viral expression before behavioral experiments began.

### Optic fiber implantation surgeries

For *in vivo* fiber photometry recording, mice were unilaterally injected with AAV1-Syn-FLEX-axon-GCaMP6s^4^ (2.2×10^13 GC/mL) in CeA (50-100 nL). For optogenetic stimulation experiments, mice received bilateral injections of AAV5-Syn-FLEX-rc[ChrimsonR-tdTomato]^5^ (2.0×10^13 GC/mL) in CeA (50-100 nL). The following coordinates for PBN/Su5 were used for both photometry (ipsilateral; 400 µm diameter core; multimode; numerical aperture (NA) 0.66; 4.0 mm length; Doric Fibers: MFC_400/430-0.66_4mm_MF1.25_FLT) and optogenetic (bilateral; 200 µm diameter core; numerical aperture (NA) 0.39; 4.0 mm length; RWD: R-FOC-L200C-39NA) fiber implantations (relative to Bregma): AP −5.2, ML ± 1.5, DV 3.5.

### GRIN lens implantation and related surgical procedures

GRIN lens implantation surgeries were performed as previously published^1^. Mice received a unilateral injection of AAV1-Syn-FLEX-axon-GCaMP6s (2.2×10^13 GC/mL) in CeA (100 nL), followed by ipsilateral implantation into the PBN of a doublet GRIN lens [GRINtech/Inscopix, NEM-050-25-10-860-DM; 0.5 mm diameter; 9.89 mm length; 250 μm focal distance on brain side at 860 nm (NA 0.47); 100 μm focal distance on air side (NA 0.19)]. Coordinates for injection into CeA and implantation of the GRIN lens above PBN are the same as for photometry coordinates described in the previous section.

### Fiber photometry recordings

Mice were habituated for two days (15 mins per day) to head-fixation with the ability to locomote on a 3D-printed circular treadmill^2^. On the third day, mice began training to lick at a spout to ingest Ensure (Ensure^®^ Plus Nutrition Shake, Vanilla, Abbott Laboratories, Chicago, IL) Company, Location) while being screened for photometry signal. For all experimental sessions, licks were detected by a capacitance-sensing lickspout (3D printed with conductive filament connected to a capacitance sensor, MPR121; Adafruit) and locomotion was recorded using an IR sensor connected to an Arduino as described in previous studies^2^.

Fiber optic cables (1 m long; 400 μm core; 0.57 NA; low autofluorescence; Doric Lenses; MFP_400/430/1100-0.57_1m_FC-MF1.25_LAF) were coupled to implanted optic fibers with zirconia sleeves (Precision Fiber Products). Black heat shrink material was placed around the fiber coupling to prevent light leakage from interfering with recordings or behavior. Excitation and emission light was passed through a fluorescence minicube allow for 405 nm and 465 nm excitation (5 port; Doric Lenses; FMC5_IE(400-410)_E(460-490)_F(500-540)_O(580-680)_S). Excitation light (~100 μW) was provided by a 405 nm or 465 nm LED (Plexon). Excitation LEDs were modulated as interleaved pulses controlled by the Nanosec photometry-behavioural system that was initially developed in a previous study^6^ (https://github.com/xzhang03/NidaqGUI).

Emission light was collected by a femtowatt photoreceiver (Newport 2151) and digitized at 2.5 kHz sampling rate (PCIe-6321; National Instruments). Data acquisition was controlled using a modified script in MATLAB (MathWorks) based on our previously developed script^6^. Further data analysis was conducted in MATLAB by first applying a least-squares linear fit to the 405 nm signal to align it to the 465 nm signal. The resulting fitted 405 nm signal was then used to normalize the 465 nm signal as follows: ΔF/F = (465 nm signal − fitted 405 nm signal)/fitted 405 nm signal. Finally, the data was z-scored using the mean and standard deviations of the baseline (pre-stimulus) segments.

Initial lick spout training sessions were used for assessing GCaMP6s signal during self-paced licking to consume Ensure. In these sessions, Ensure delivery was conditional on a lick detected on the lick spout and Ensure deliveries could be triggered as often as every 5 s. More specifically, the first 2 s served as the lick window for triggering Ensure delivery, and the last 3 s served as a time out window to allow the mouse to finish consuming the bolus of Ensure before another delivery was triggered. This timing was chosen as it openly results in continuous binge-lick consumption by mice based on our prior studies^1^. Ensure delivery was available for the first 20 mins and photometry recording continued for an additional 5 mins after the session’s end to capture lick bouts in the absence of Ensure consumption for analysis. Mice were food restricted prior to these training sessions (see above in Experimental model and subject details).

### Pavlovian conditioning, operant conditioning, taste, and tail shock procedures

To condition licking behavior, food restricted mice were first trained for 3-5 days on a Pavlovian protocol in which a 5 s blue light cue was presented followed by a 50 ms delay after the cue offset, then delivery of a bolus of Ensure to the lick spout. After this initial Pavlovian conditioning, mice moved on to 3-4 days of an operant lick-conditional protocol in which Ensure delivery was conditional on licking during the cue period. In this protocol, if at least one lick was detected during the 5 s cue period, Ensure was delivered to the lick spout 50 ms after cue offset, but if the mouse did not lick during the cue period, then no Ensure was delivered. For both phases of conditioning, sessions were ~12 min in duration, with only one session per day, each consisting of 25 trials with a variable inter-trial interval of 30 s.

To determine if axon activity was different as a result of different tastants, subsequent operant sessions replaced Ensure with water (2-3 sessions per mouse), 1 mM quinine (1 session), and 5%, 10%, or 20% sucrose (3 sessions per mouse for each concentration of sucrose). For water and quinine, mice were water restriction prior to experimental sessions (see above), and for sucrose sessions mice were food restricted prior.

We tested responses to tail shock as previously described^2^. Briefly, mice were first habituated to tail shock pads adhered to the tail for 15 mins per day for two days. On each of 3 tail shock test days, mice received a 0.3 mA tail shock stimulus (6 × 50 ms duration tail shock train, 6 Hz) every 30 s for the 10 min session.

### Two-photon imaging procedures

Two-photon imaging was performed using a resonant-scanning two-photon microscope (Thorlabs Bergamo) at ~31 frames s^−1^ and 512 × 512 pixels per frame. Excitation, 960 nm, was achieved using a femtosecond tunable laser (Coherent Discovery NX). Imaging was performed with a 4x 0.2 NA air objective (Nikon) that matches the numerical aperture of the GRIN lens. Imaging fields of view were at a depth of 100–300 μm below the face of the GRIN lens. Laser power ranged between 40-60 mW at the front aperture of the objective (the power at the sample was substantially less because of partial transmission via the GRIN lens). Head-fixation, self-paced licking to consume Ensure, and tail shock habituation and testing procedures are same as above.

### Optogenetic stimulation

For all optogenetic experiments, mice underwent patch cord (1 m long, 200 μm core, 0.39 NA; RWD), habituation for 15-30 mins per day for 2-3 days prior to the start of experimental sessions. Arduino-controlled drivers mediated the photostimulation from either a 620 nm LED (Plexon) or a 635 nm laser (Shanghai Laser) for head-fixed and freely moving experiments, respectively. All stimulation was bilateral unless otherwise specified in the legends. Stimulation was always 10 ms pulses with 5 mW power at the specified frequencies.

### Head-fixed feeding experiments with optogenetic stimulation

We used a head-fixed behavioral apparatus across several optogenetic experiment protocols to achieve precise temporal resolution for assessing the effects of photostimulation on orofacial movements and ingestion. Following habituation sessions, food restricted mice were trained without photostimulation for 3 days (30 mins per day) to lick at a lick spout to receive Ensure (see *self-paced licking to consume Ensure* described above in fiber photometry recordings). In testing sessions with photostimulation (5 mW, 10 ms pulses), mice underwent 3 consecutive runs while licking for Ensure: 5 mins of no stimulation, 5 mins with the stimulation protocol chosen for that session, 5 mins of no stimulation. In these sessions, we examined the effects of four different stimulation protocols. For the lick bout-triggered photostimulation protocol, stimulation was delivered once 2 licks occurring <1 s apart was detected and ended when an interlick interval of >1 s occurred or 10 s had elapsed, whichever occurred first. This protocol was tested across multiple stimulation frequencies: 0 Hz, 5 Hz, 10 Hz, 15 Hz, and 20 Hz. After determining that the 20 Hz frequency stimulation induced the greatest behavioral effect, 20 Hz was used for the remainder of the optogenetic experiments. The other stimulation protocols included continuous photostimulation (20 Hz for the entire 5 min stimulation run), lick-triggered Ensure delivery with 3 s photostimulation (stimulation occurred simultaneously with Ensure delivery and lasted for 3 s), and lick-triggered photostimulation (a single pulse administered each time a lick was detected). Mice in these experiments performed each stimulation protocol for 2-3 sessions. For control sessions, all 3 runs were conducted without photostimulation, and the second run was used for the “No Stim” data to compare against the various stimulation protocols.

To assess the effect of photostimulation-induced ingestion of an aversive substance, *self-paced licking to consume* sessions were also run using 1 mM quinine (2-3 sessions per mouse) in place of Ensure under the continuous photostimulation protocol (5 mW, 20 Hz). For this experiment, mice were *ad lib*. fed but water restricted.

To test the interaction between photostimulation and the influences of satiation and licking fatigue, longer sessions were conducted using *ad lib*. fed mice. These sessions lasted 45 min with continuous 20 Hz stimulation. For investigating the effects of satiation on lick microstructure during photostimulation, the amount of Ensure dispensed was recorded every 5 mins in addition to quantification of detected licks. For isolating the influence of licking fatigue, the lick spout was available but no Ensure was delivered throughout the session.

In order to assess the effects of photostimulation prior to any experience with the lick spout, a new cohort of mice was prepared. These *ad lib*. fed mice underwent head-fixation and patch cord habituation, but their first test session was also their first exposure to the lick spout (Naïve session, no Ensure deliveries) prior to any lick spout training. They then trained for 1-2 days to lick at the lick spout for Ensure (see above: *self-paced licking to consume Ensure*), followed by an additional 4-12 training sessions with stimulation runs and, finally, the session used as “Experienced” data. Paw-to-mouth behaviors were manually scored by a trained observer from videography of experiment sessions (video acquisition described below). Locomotion and lick spout data were collected for all head-fix sessions utilizing the running wheel and lick spout (same as described above in fiber photometry recordings).

For the pellet grasping task, the head-fixed apparatus was modified based on a prior study^7^ in order to train mice to reach for food pellets. The running wheel was replaced by a raised platform with an affixed 3D-printed hut for the mouse to sit in. A rod was secured to the front of the hut at elbow-level for the mouse to rest its forepaws while not eating. Food restricted mice were habituated for 2-3 sessions to the hut. After habituation, an experimenter began manually delivering grain pellets (5TUM, 1811143, TestDiet, Richmond, IN) to the mouse’s mouth using forceps. Pellets were offered at progressively increasing distance from the mouse’s mouth across the course of the 2-4 day training period. Once the mouse consistently retrieved pellets with its tongue and assumed a comfortable bipedal posture while eating, a pellet dispenser replaced the experimenter in delivering the grain pellets. The dispenser was placed at elbow-height and angled towards the mouse’s preferred forepaw, and the mouse was trained for another 3-8 days to reach for, grasp, and consume the pellets from the dispenser. During experimental sessions, mice were given continuous access to pellets via the dispenser for 10 mins without stimulation, then for 10 mins with stimulation (5 mW, 20 Hz) and the number of pellets successfully grasped and consumed was manually quantified.

For all head-fixed experiments, a Thorlabs Zelux 1.3MP camera acquired videography at 59-62 frames per second (fps) of the mouse performing licking and other orofacial behaviors during testing sessions.

### Freely moving food and water intake experiments with optogenetic stimulation

24 hours before a freely moving food restricted feeding test session, mice were placed in a clean cage with fresh bedding and without food. For mice tested after being on *ad lib*. chow, they remained in their home cage until just before the session. Across all freely moving testing parameters, mice were briefly head-fixed for patch cord tethering, then placed in the cage where testing would take place. At the start of the 2 h freely moving feeding assay, mice were provided a single 3 g standard chow pellet. The pellet was weighed every 30 mins for the 2 h assay, during which time the mice either received continuous photostimulation (5 mW, 20 Hz) for the first hour and no stimulation for the second hour or underwent control sessions without stimulation for the duration of the 2 h feeding assay. Freely moving feeding assay sessions testing the effects of different stimulation frequencies (5 mW; 0 Hz, 5 Hz, 10 Hz, 15 Hz, and 20 Hz) employed the same behavioral protocol as just described.

For experiments investigating the interacting effect of food proximity on photostimulation-induced consumption, after patch cord tethering, *ad lib*. fed mice were placed in a new cage without bedding and provided a single ~3 g standard chow pellet. The 30 mins assay was broken into 3 time periods: 10 mins of no stimulation, 10 mins with experimenter-delivered stimulation (5 mW, 20 Hz), and a final 10 mins no stimulation period. In the *stim near food* protocol, during the second 10 min period of the session, a trained experimenter administered photostimulation whenever the mouse approached the chow pellet and halted the stimulation when the mouse distanced themselves from the pellet. For the *stim away from food* protocol, the experimenter only delivered photostimulation when the mouse was not near the chow pellet (i.e., in another area of the cage and not within reach of the food) during the 10 min stimulation period of the assay. For both session types, the chow pellet was weighed every 10 mins for the 30 mins assay.

We also conducted a 2 h feeding assay using the *stim near food* protocol to examine the impact of this proximity-based stimulation across a longer time course and evaluate satiation effects. Just as described above, a trained experimenter turned on the photostimulation whenever the mouse was near the chow pellet but did so throughout the entire 2 h assay. *Ad lib*. fed mice were used in these experiments and the chow pellet was weighed every 15 mins. The control sessions for this assay were 2 h feeding sessions (chow pellet weighed every 30 mins) where no photostimulation was delivered at any point.

For freely moving water intake experiments, mice were water restricted (described above) and habituated to drinking from a 35mm sized Petri dish. There were 2-4 habituation sessions lasting 30 mins-1 h in duration that took place in a clean, bedding-less cage where the Petri dish filled with water was affixed to the cage floor using adhesive mounting putty to prevent spilling. Each mouse performed 2-3 sessions each for the optogenetic stimulation condition (5 mW, 20 Hz continuous photostimulation for the first 30 mins and no stimulation for the second 30 mins) and the control condition (no stimulation sessions). An experimenter carefully removed and weighed the Petri dish every 15 mins with the intent to measure water intake, but we discovered that the mice were primarily chewing and consuming the plastic dish instead of drinking the water from it.

Videography of these experiments were captured (AMcap) from a camera mounted on the side and from the top (Arducam). Videos were acquired at 30 fps, and video-based tracking of the mouse and food pellet was performed using DeepLabCut (detailed in separate section). To generate the DeepLabCut model, the mouse’s nose and the food pellet were manually labeled in an average of 135 frames per video extracted from 4 videos captured from above. Additionally, the optic fibers were labeled only in frames where the laser was on, allowing for analysis of the duration of stimulation during closed-loop sessions. Videography from these sessions were manually scored by a trained observer for quantifying and analyzing cage licking and fictive eating (i.e. paws raise to mouth as if eating a non-existent pellet) behaviors.

### DeepLabCut & SimBA model training

For markerless tracking of the jaw and tongue, a DeepLabCut^8^ network was trained using a ResNet-50 backbone. A training dataset was prepared by extracting an average of 215 frames per video from 53 videos, captured under varied lighting conditions, to manually label the two anatomical landmarks of interest. If the tongue was not visible, the inside of the mouth where the tongue would naturally rest was labeled. The data sets for each head-fixed and freely moving optogenetic experiments were shuffled and split 95:5 for training and testing, respectively. Frames with poor tracking were extracted and corrected using DeepLabCut’s refinement GUI, and the model was retrained until satisfactory tracking results were obtained. Using a confidence threshold of 0.9, the final model for head-fixed experiments achieved a test error of 2.28 pixels and a train error of 2.14 pixels compared to human-provided annotations, and a test error of 3.6 pixels and a train error of 2.14 pixels for freely moving experiments. For head-fixed experiments, tracking data was subsequently imported into the Simple Behavioral Analysis toolkit^9^, where a model was trained to identify licks using a random forest approach. Approximately 5,300 frames from 11 videos were selected for training and manually annotated within SimBA as containing or not containing a lick. For all experiments analyzed with DeepLabCut, pixel coordinates were converted to millimeters using SimBA’s video parameters GUI.

### Optogenetic stimulation with toy bug

To determine if photostimulating this pathway could induce hunting-like behaviors towards an artificial prey, we adapted a protocol from a previous study^10^. Mice, *ad lib*. fed, were briefly head-fixed for patch cable tethering and placed in an empty, bedding-less cage. The robotic cricket (HEXBUG nano flash) was released in an unoccupied spaced in the cage and continuous stimulation (5 mW, 20 Hz) was delivered for 5 mins before terminating and the artificial prey was removed. A trained observer manually quantified the number of bites and bite duration from videography (AMcap) of these sessions obtained from a camera mounted on the side and from the top (Arducam) of the cage acquired at 30 fps.

### Real-time place preference assay

Mice underwent 2 days of habituation (one 20 min session / day) to a rectangular arena (510 mm long, 270 mm wide, 300 mm tall) that was divided in the middle by a partial wall. An overhead camera (Microsoft USB webcam) and mouse tracking software (EthoVision; Version 14) tracked animal movement and mediated closed-loop photostimulation for the real-time place preference assay. Across three testing days, mice were placed in arena for 10 mins with no photostimulation. Following this baseline period, the left side of the arena was designated as the active side, meaning photostimulation was delivered whenever the entirety of the mouse was present on the left side (5 mW, 20 Hz). Locomotion speed and time spent on each side of the arena was recorded. Videography from these sessions were manually scored by a trained observer for quantifying and analyzing chewing behaviors.

### Histology

Mice were transcardially perfused with fresh 4% paraformaldehyde (PFA). For fiber photometry and two-photon experiments, brains were immediately extracted. For optogenetic experiments, perfused mice were decapitated and heads were left in 4% PFA with headposts and optic fibers left in place for up to a week. After extraction, brains were left in fresh 4% PFA overnight, cryoprotected with 20% sucrose for a minimum of 48 h, the cut into 3 series of 40 μm free-floating sections with a Leica VT1200S vibratome. Sections were mounted onto slides and coverslipped using VECTASHIELD® Vibrance™ Antifade Mounting Medium with DAPI (Vector Labs). Sections were imaged at 20 x magnification using a digital slide scanner (VS200, Olympus, Japan). All viral expression and optic fiber and GRIN lens implant placements were verified by post hoc histology using “The Mouse Brain in Stereotaxic Coordinates” by Paxinos and Franklin (2^nd^ edition, 2001). Mice with incorrect virus expression location or implant placement were excluded from the study.

### QUANTIFICATION AND STATISTICAL ANALYSIS

Photometry and two photon imaging data were first processed in MATLAB. All statistical tests were performed in GraphPad Prism 10. Statistical details and sample sizes are found in the figure legends. Normality tests were first performed to determine the appropriate statistical test to use. No statistical methods were used to pre-determine sample sizes, but our sample sizes were chosen to reliably measure experimental parameters while remaining in compliance with ethical guidelines for minimizing animal use and were similar to those reported in previous publications. Significance levels are indicated as follows: *p<0.05, **p<0.01, ***p<0.001, ****p<0.0001. All data are presented as mean ± S.E.M. Figures were generated in MATLAB or GraphPad Prism 10 and exported to Adobe Illustrator.

## SUPPLEMENTAL MOVIE DESCRIPTIONS

**Supplemental Movie 1: Tracking of tongue and jaw positions from videography using DeepLabCut analysis**

Beginning of movie shows side-by-side views of the same mouse either with no photostimulation (left) or receiving 20 Hz continuous photostimulation (right). The second half of the movie shows the effect of photostimulation on the jaw trajectory from periods of the movie in which no licking is occurring.

**Supplemental Movie 2: Example of pellet grasping behavior in head-fixed mice with and without photostimulation of CeA**^**pons**^ **axons**

Movie shows side-by-side views of the same *ad lib*. fed mouse either with no photostimulation (left) or receiving 20 Hz continuous photostimulation (right).

**Supplemental Movie 3: Example of fictive eating behaviors overriding normal pellet ingestion in a food restricted mouse**

Movie shows a food restricted mouse either receiving 20 Hz continuous photostimulation and performing fictive eating behaviors instead of eating the available food pellet.

**Supplemental Movie 4: Example of CeA**^**pons**^ **photostimulation causing biting of a toy bug**

Movie top and side views of an example *ad lib*. fed mouse exposed to a toy bug (artificial prey) with no stimulation followed by with 20 Hz continuous photostimulation.

**Supplemental Movie 5: Examples of closed-loop CeA**^**pons**^ **photostimulation driving food pellet grasping and biting in freely moving mice**

Movie shows two examples of an *ad lib*. fed mouse receiving 20 Hz photostimulation only when it approaches and is near food. This stimulation leads to lunging, grasping, biting, and retrieving of the food pellet.

**Supplemental Movie 6: Closed-loop CeA**^**pons**^ **photostimulation either when the mouse is near food or when it is away from food**

Beginning of the movie shows an *ad lib*. fed mouse either receiving 20 Hz photostimulation when the mouse is near food (top) or away from food (bottom). The second part of the movie shows top and side views of an example *ad lib*. fed mouse receiving 20 Hz photostimulation when it is near food for 2 hours (movie is sped by binning to ~3 frames per minute).

**Supplemental Movie 7: Examples of fictive eating and cage licking behavior observed during CeA**^**pons**^ **photostimulation when the mouse is away from food**

Movie shows two examples of fictive eating followed by two examples of cage licking from *ad lib*. fed mice receiving 20 Hz photostimulation when away from food.

**Supplemental Figure 1.**
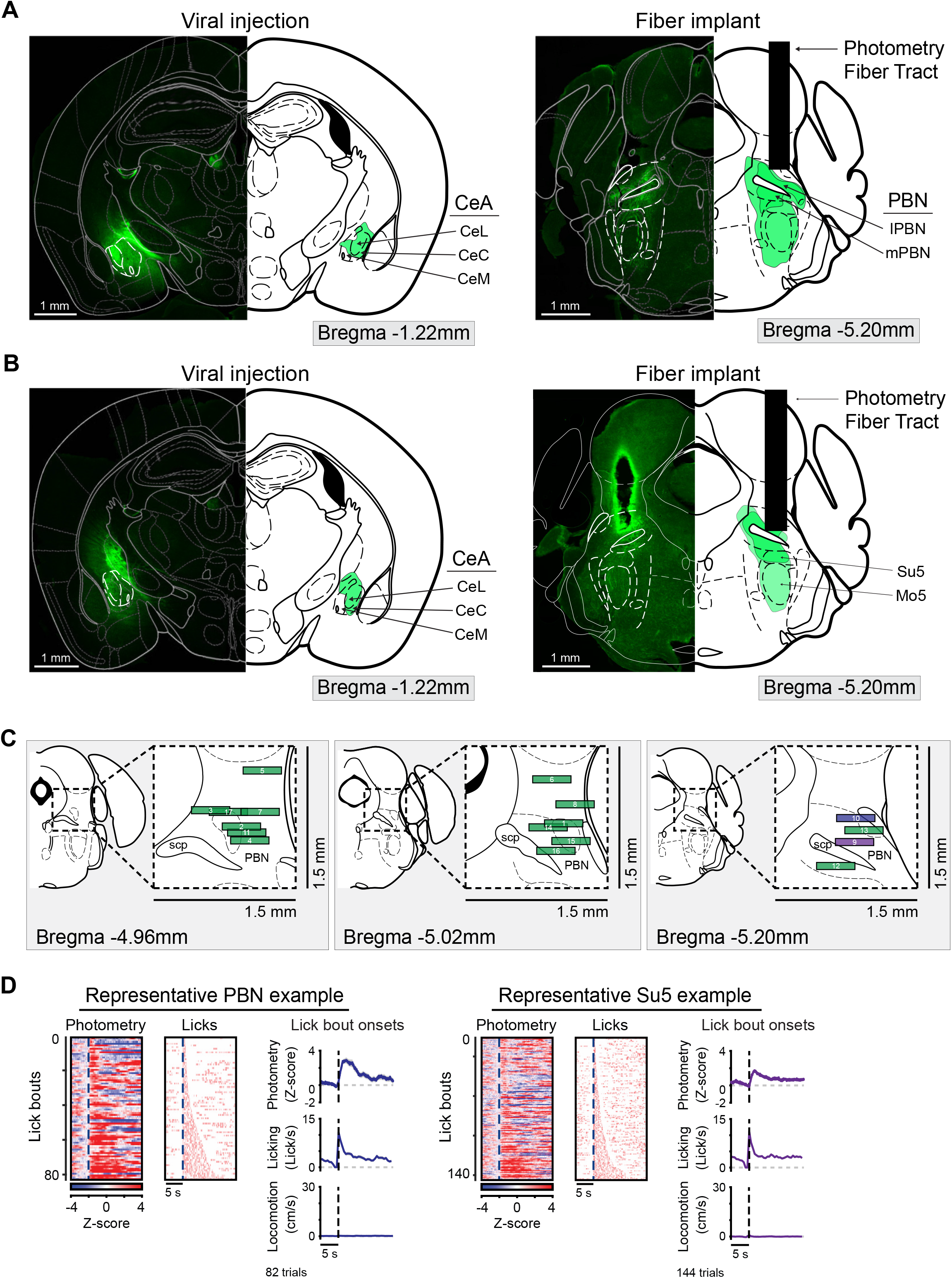
Histology of photometry mice and comparison of signals from probable PBN versus Su5 targets. (A) Left: axon-GCaMP6s expression at the injection site in the CeA. Right: axon-GCaMP6s expressing in axons within dorsolateral pons below the implanted photometry fiber, which was located above the PBN. (B) Left: axon-GCaMP6s expression at the injection site in the CeA. Right: axon-GCaMP6s expressing in axons within dorsolateral pons below the implanted photometry fiber, which end up implanted through dorsal PBN and likely pick up axonal signals from within mPBN and Su5. (C) Locations of implanted fibers based on histology from all mice used in photometry experiments. (D) Left: heatmaps and time courses of photometry, licking and locomotion from a mouse with the fiber implanted above the PBN. Right: heatmaps and time courses of photometry, licking and locomotion from a mouse with the fiber implanted into the PBN and above the Su5.

**Supplemental Figure 2.**
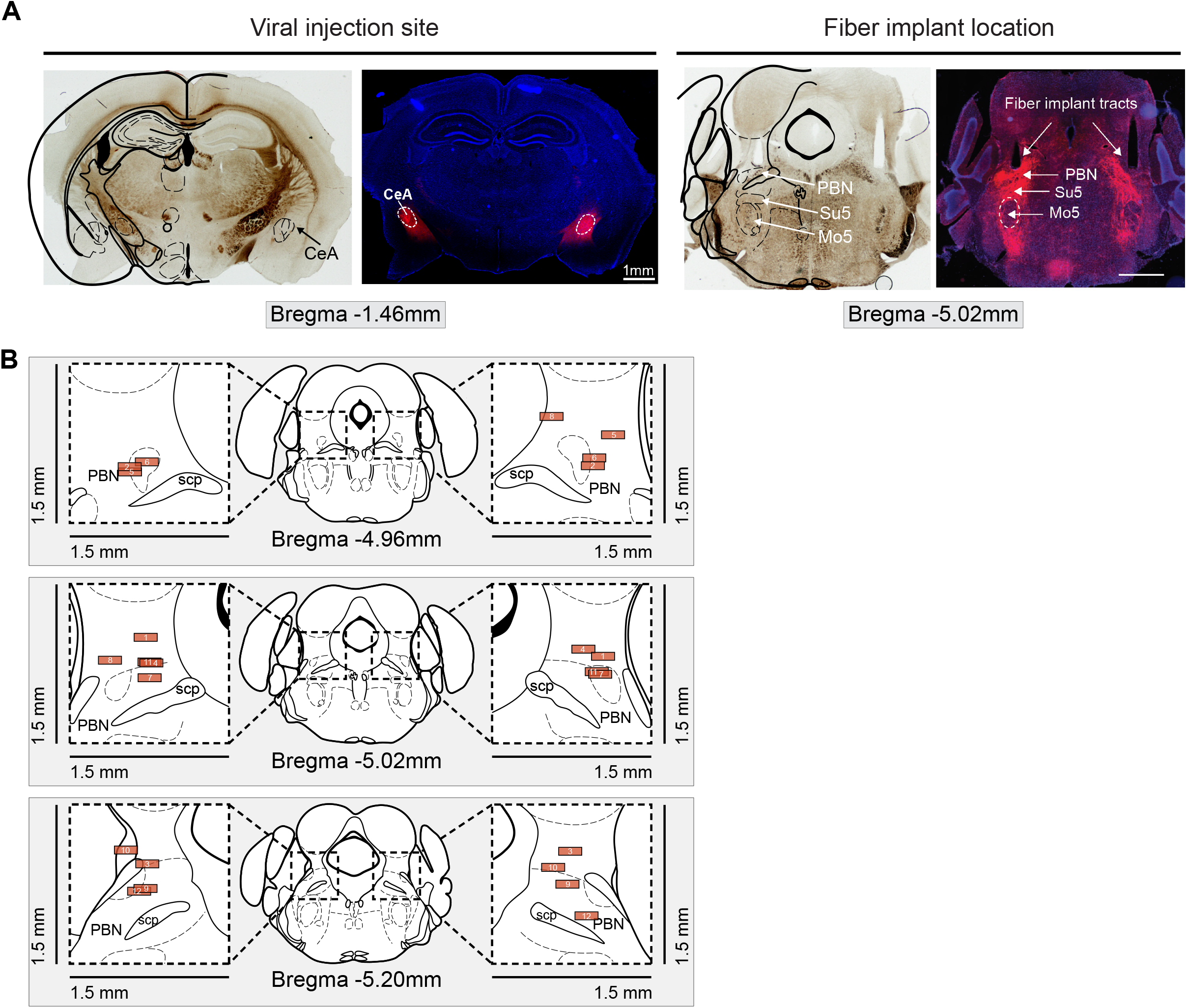
Histology of mice used in the optogenetic studies. (A) Left: Representative brightfield and epifluorescence images depicting bilateral ChrimsonR-tdTomato expression at the CeA injection sites. Right: ChrimsonR-tdTomato-expressing axon terminals in the pons subregions located below the implanted optogenetic fibers. (B) Locations of implanted fibers based on histology from all mice used in optogenetic experiments.

**Supplemental Figure 3.**
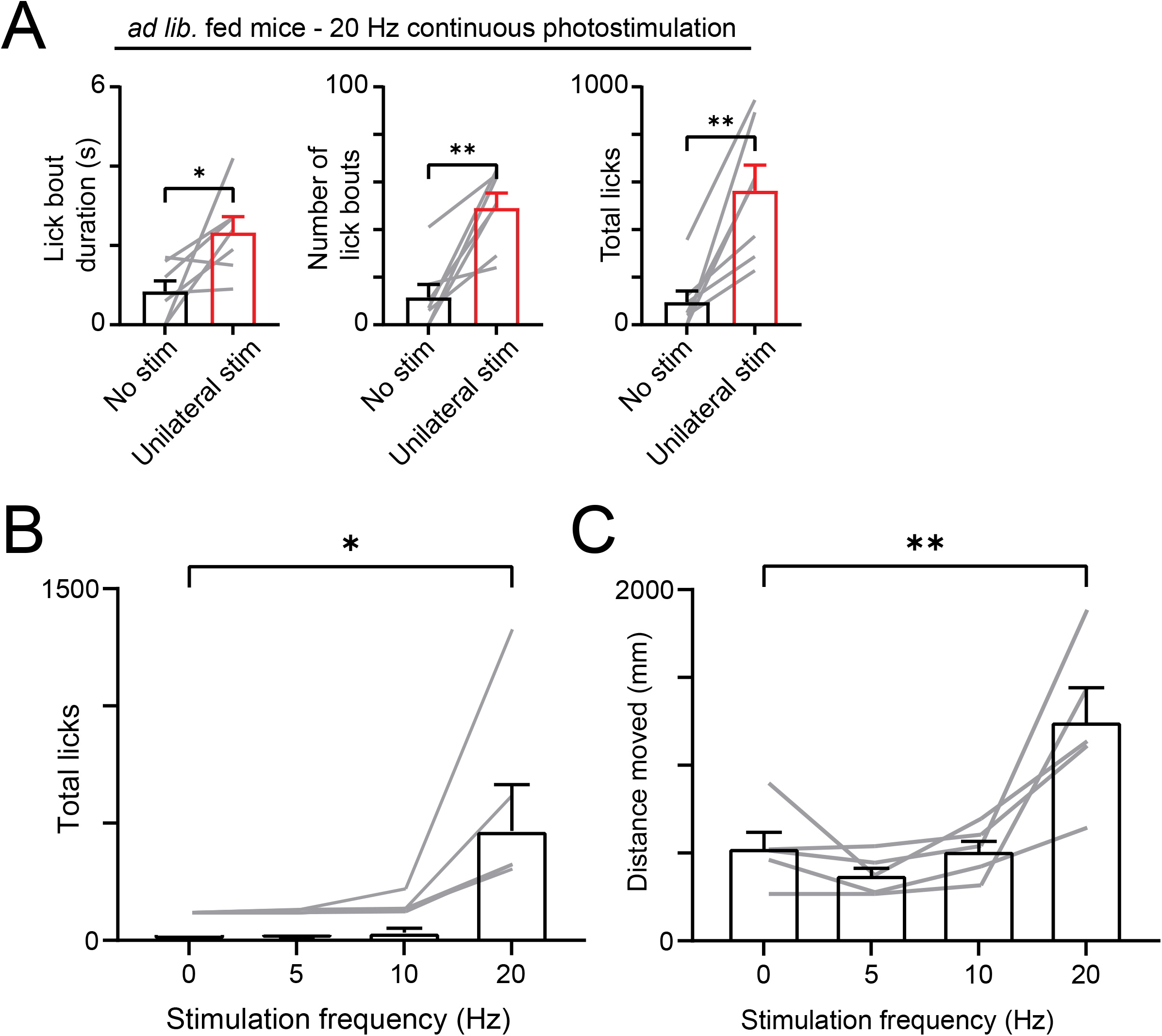
Additional analyses of head-fixed optogenetic experiments. (A) The effect of unilateral 20 Hz continuous stimulation on lick bout duration (left), number of lick bouts initiated (middle), and total number of licks (right) in *ad lib*. fed mice. Two-tailed paired t-test. N = 7 mice (3M, 4F). (B) Analysis of licks detected by videography in *ad lib*. fed mice during photostimulation of varying frequency. Repeated-measures one-way ANOVA with Dunnett’s correction for multiple comparisons. N = 5 mice (3M, 2F). (C) Analysis of jaw movement distance measured by videography in *ad lib*. fed mice during photostimulation of varying frequency. Jaw movement related to tongue extension is not included in this analysis. Repeated-measures one-way ANOVA with Dunnett’s correction for multiple comparisons. N = 5 mice (3M, 2F).

**Supplemental Figure 4.**
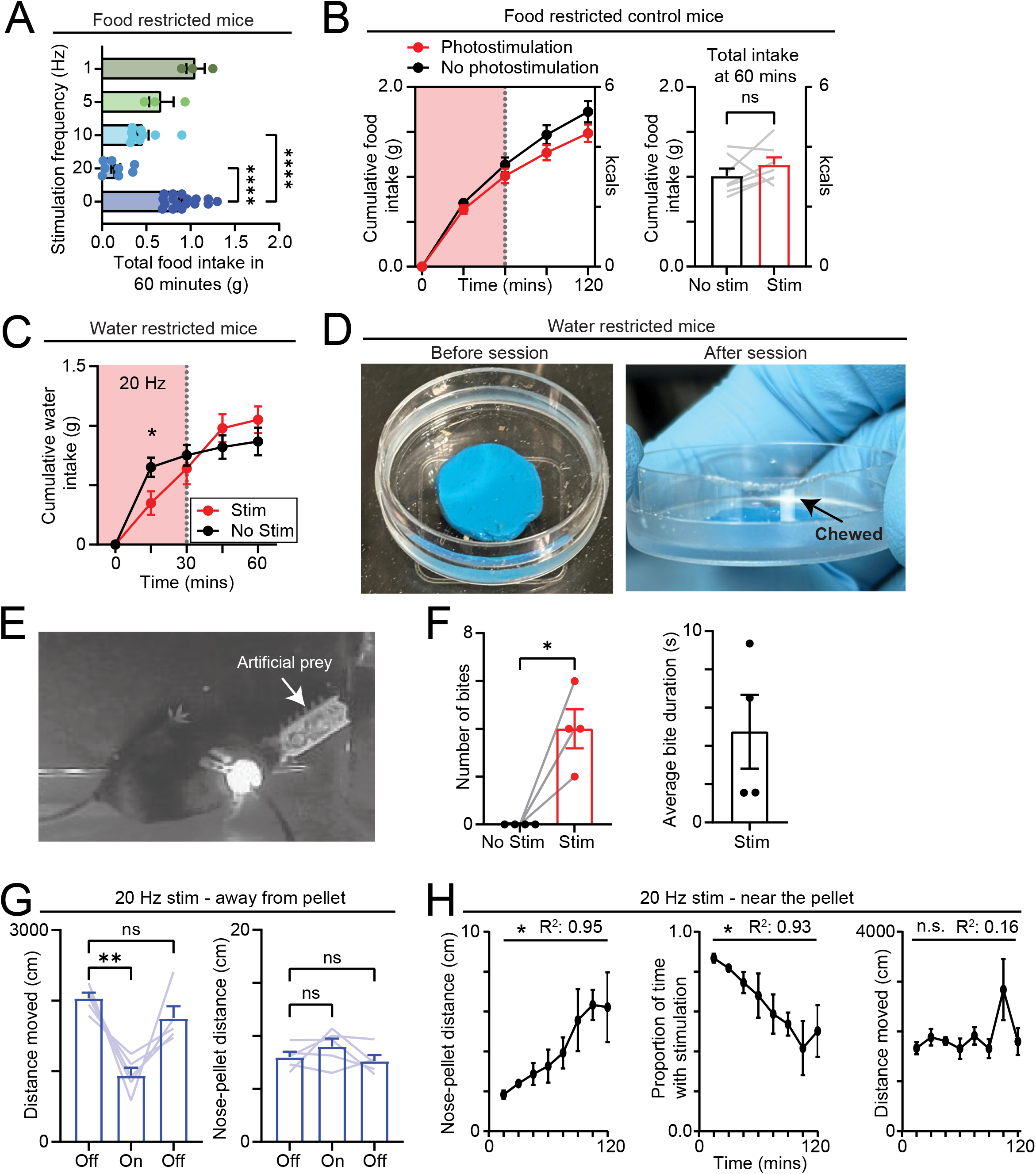
Additional analyses of freely moving optogenetic experiments. (A) Total food consumed in the first 60 minutes without or with photostimulation at different frequencies. Mixed-effects model. N = 17 mice (9M, 8F) without photostimulation, N = 3 mice (3M) with 1 Hz, N = 3 mice (3M) with 5 Hz, N = 9 mice (6M, 3F) with 10 Hz, N = 8 mice (3M, 5F) with 20 Hz. (B) Left: time course of average food intake of food restricted negative control mice (no Chrimson expression) over 2 hours with either 20 Hz continuous or no photostimulation during the first 60 minutes. Right: total food consumed in the first 60 minutes with or without photostimulation. Two-tailed paired t-test. N = 7 mice (4M, 3F). (C) Time course of average water intake of water restricted mice over 1 hours with either 20 Hz continuous or no photostimulation during the first 30 minutes. Repeated-measures two-way ANOVA with Šidák’s correction for multiple comparisons. N = 5 mice (3M, 2F). (D) Images depicting 30 mm plastic petri dish used for water intake experiments before and after stimulation from a single example session. We often observed chewing of the petri dish and likely ingestion of the plastic as no plastic debris on cage floor was observed. (E) Image depicting a mouse biting an artificial prey (toy bug) while receiving 20 Hz photostimulation. (F) Quantification of the number of bites onto the artificial prey and average bit duration during 20 Hz photostimulation. Two-tailed paired t-test. N = 4 mice (2M, 2F). (G) Left: distance moved by mice during 20 Hz photostimulation that was triggered if mice were not nearby the food pellet. During the 10 minutes of stimulation (On), mice moved less and were more engaged in fictive eating behaviors. Right: the distance between the nose and the pellet were not significantly different during On or Off stimulation periods. (H) Left: the distance between the nose and the pellet over the course of 2 hours of close-loop stimulation (20 Hz) just when the mouse was near the food pellet. There was a significant correlation between the nose-pellet distance and time. Middle: the proportion of time during each 15-minute bin that close-loop stimulation (20 Hz) was active. There was a significant correlation between the proportion of time with stimulation and overall time. Right: the distance moved during each 15-minute bin was not correlated with time.

**Supplemental Figure 5.**
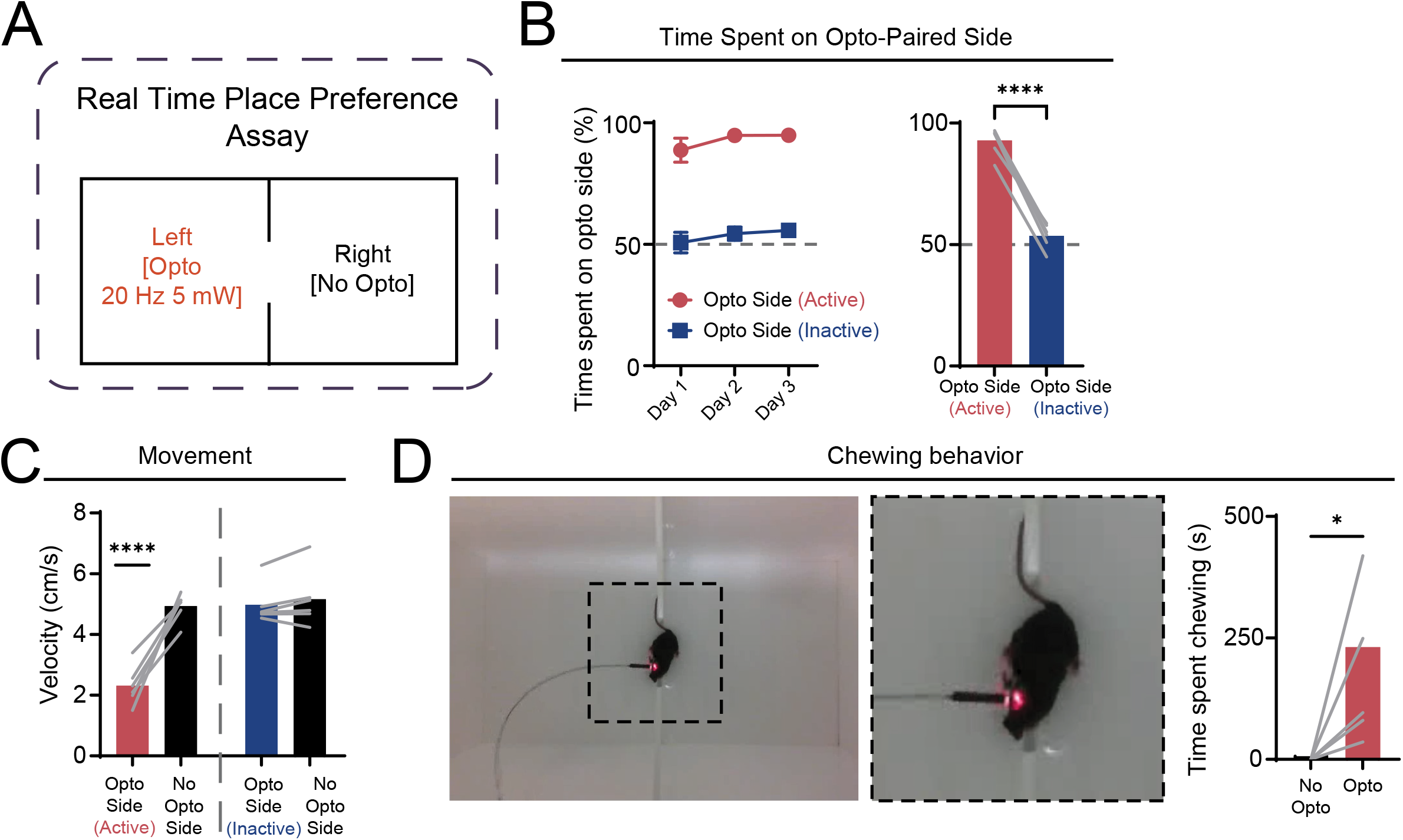
Real-time place preference assay. (A) Schematic of the real-time place preference chamber in which the left side of the chamber delivered continuous 20 Hz (635 nm laser; 5 mW) if the mouse entered that side and persisted until the mouse exited that side of the chamber. Each day involved a 10-minute exploration period during which no photostimulation was delivered on either side of the chamber followed by the 20-minute experiment in which the left side did deliver photostimulation. (B) Percentage of time spent on the left side of the chamber during 10-minute exploration period in which stimulation was not activated (blue) and the 20-minute period in which photostimulation was active (red). One-sample t-test with a theoretical mean of 50 %. Right: Comparison of the average percentage of time spent on the opto side when the photostimulation was active versus when it was inactive. Two-sample paired t-test. (C) Movement velocity on each side of the chamber during sessions in which the opto side was active or inactive. Mixed-effects analysis with Tukey’s correction for multiple comparisons. All data: N = 6 mice (4M, 2F).

## Notes

### Competing Interest Statement

The authors have declared no competing interest.

